# Gene replacement of α-globin with β-globin restores hemoglobin balance in β-thalassemia-derived hematopoietic stem and progenitor cells

**DOI:** 10.1101/2020.10.28.359315

**Authors:** M. Kyle Cromer, Joab Camarena, Renata M. Martin, Benjamin J. Lesch, Christopher A. Vakulskas, Viktor T. Lemgart, Yankai Zhang, Ankush Goyal, Feifei Zhao, Ezequiel Ponce, Wai Srifa, Rasmus O. Bak, Naoya Uchida, Ravindra Majeti, Vivien A. Sheehan, John F. Tisdale, Daniel P. Dever, Matthew H. Porteus

## Abstract

β-thalassemia pathology is not only due to loss of β-globin (*HBB*), but also erythrotoxic accumulation and aggregation of the β-globin binding partner, α-globin (*HBA1/2*). Here we describe a Cas9/AAV6-mediated genome editing strategy that can replace the entire *HBA1* gene with a full-length *HBB* transgene in β-thalassemia-derived hematopoietic stem and progenitor cells (HSPCs), which is sufficient to normalize β-globin:α-globin mRNA and protein ratios and restore functional adult hemoglobin tetramers in patient-derived red blood cells. Edited HSPCs were capable of long-term and bi-lineage hematopoietic reconstitution in mice, establishing proof-of-concept for replacement of *HBA1* with *HBB* as a novel therapeutic strategy for curing β-thalassemia.

---

β-thalassemia is one of the most common genetic blood disorders in the world, with a global annual incidence of 1 in 100,000^1^. Patients with this disease suffer from severe hemolytic anemia and, even with intensive medical care, experience a median life expectancy of approximately 30 years of age^2,3^. The most severe form of the disease—β-thalassemia major—is caused by homozygous (or compound heterozygous) loss-of-function mutations throughout the β-globin (*HBB*) gene. This results in loss of HBB protein, reducing the ability of red blood cells (RBCs) to deliver oxygen throughout the body, as well as accumulation of unpaired α-globin (from *HBA1* and *HBA2* genes), leading to dramatic erythrotoxicity and ultimately hemolysis. In fact, disease severity is known to be directly correlated with the degree of imbalance between β-globin and α-globin chains^4^. The current standard of care for β-thalassemia involves frequent blood transfusions and/or iron chelation therapy, making this one of the most costly genetic diseases in young adults^5^. Currently, the only curative strategy for this disease is allogeneic hematopoietic stem cell transplantation (HSCT) from an immunologically matched donor. However, in the majority of cases, no matched donor is available for allogeneic HSCT, and, even if one is identified, transplants from these donors carry a high risk of immune rejection and graft-versus-host disease^6^.

Therefore, an ideal treatment would involve isolation of patient-derived hematopoietic stem and progenitor cells (HSPCs), introduction of *HBB* to restore HBB protein levels, followed by autologous HSCT of the patient’s *own* corrected HSPCs that carries no risk of immune rejection. Employing this logic, several gene therapies have been developed as a potentially curative measure for β-thalassemia, primarily through delivery of an *HBB* transgene using lentiviral vectors^7–9^. While these approaches have been shown to restore HBB to therapeutic levels in human clinical trials for β-thalassemia^10^, in one case the therapeutic benefit was achieved from a single dominant clone^11^. In addition to this, while no adverse events have occurred after a several-year follow up, integrations into tumor suppressors such as *NF1* are commonly observed, leading to safety concerns regarding semi-random integration of transgenes into the genome^12^.

Therefore, alternative strategies that rely on the reported safety^13,14^ of more precise genomic changes are being developed, such as CRISPR/Cas9-mediated inactivation of a repressor of fetal hemoglobin, the upregulation of which could compensate for the lack of HBB^15^. However, there is some concern with this approach that the upregulation of fetal hemoglobin may not be sufficient to rescue the β-globin:α-globin imbalance and that the upregulation may not persist in adult patients in whom fetal hemoglobin is naturally silenced^16,17^. Moreover, this approach does not address the genetic cause of β-thalassemia—inactivation of *HBB*—and may not sufficiently rescue the disease phenotype *in vivo*. Furthermore, all of these therapies act to compensate *only* for the lack of HBB, and do not diminish levels of α-globin. This led us to explore whether Cas9/AAV6-mediated genome editing could be used to address both molecular factors responsible for disease pathology in a patient’s own HSPCs, which may be an effective and safe strategy to restore the balance between β-globin and α-globin and ultimately correct β-thalassemia.

In this study, we have leveraged the combined Cas9/AAV6 genome editing method to mediate site-specific gene replacement of the endogenous *HBA1* gene with a full-length *HBB* transgene while leaving the highly-homologous *HBA2* gene unperturbed. We found that this process allowed us to replace the entire coding region of *HBA1* with an *HBB* transgene at high frequencies, which both normalized the β-globin:α-globin imbalance in β-thalassemia-derived HSPCs and rescued functional adult hemoglobin tetramers in RBCs. Following transplantation experiments into immunodeficient NSG mice, we found that edited HSCs were able to repopulate the hematopoietic system *in vivo* and could potentiate long-term engraftment, indicating that the editing process does not disrupt normal hematopoietic stem cell function.

To our knowledge, this is the first time that a single Cas9-induced DSB has been shown to mediate replacement of an entire endogenous gene with an exogenous transgene at high frequencies. Because of this, we expect our findings to be broadly applicable to a wide variety of monogenic diseases caused by loss-of-function mutations scattered throughout a particular gene, ultimately expanding the genome editing toolbox.

## RESULTS

Cas9/AAV6-mediated genome editing is a robust system capable of introducing large genomic integrations at high frequencies across many loci in a wide variety of cell types, including HSPCs^18–23^. In fact, CRISPR-mediated approaches have been successfully employed to correct the disease-causing SNPs at high frequencies in HSPCs^22–27^. However, β-thalassemia is caused by loss-of-function mutations scattered throughout the gene, rather than the single polymorphism responsible for SCD. Therefore, a universal correction scheme for all patients requires delivery of a full-length copy of *HBB*. The simplest method for doing so would be to knock in a functional *HBB* transgene at the endogenous locus. However, this approach suffers from a number of technical issues: 1) Cas9-mediated DSBs in *HBB* could disrupt partially-functional alleles in β-thalassemia minor and intermediate patients; 2) codon divergence is required to introduce a fulllength β-globin cDNA at the endogenous locus in order to prevent partial recombination surrounding the Cas9 DSB site, which can negatively affect transgene expression levels^28^; 3) introns must be removed since they cannot be rationally diverged, which may disrupt the important functional role that the *HBB* introns are known to play in gene regulation^29,30^; 4) disease-causing mutations in surrounding regulatory regions are likely to persist after introduction of diverged cDNA; and 5) this strategy would be ineffective for many patients with disease caused by large deletions of the β-globin locus **(Supplemental Fig. 1)**^31^.

Prior work has shown that β-thalassemia patients with lowered α-globin levels demonstrate a less severe disease phenotype^4,32^. Therefore, knock-in of full-length *HBB* at the α-globin locus could most effectively allow us to improve the β-globin:α-globin imbalance in a single genome editing event while overcoming the problems inherent with introducing *HBB* into the endogenous locus.

### Efficient and specific indel formation in the highly homologous α-globin genes

Because α-globin is expressed from two genes (*HBA1* and *HBA2*) as HSPCs differentiate into RBCs, we hypothesized that site-specific replacement of a single α-globin gene with *HBB* could allow us to achieve RBC-specific expression of *HBB* without eliminating critical α-globin production. Though the *HBA1* and *HBA2* genes are extremely homologous (5’ UTR, all three exons, and intron 1 are 100% homologous; intron 2 is 94.0% homologous; 3’ UTR is 83.8% homologous), we were able to identify a limited number of Cas9 single guide RNA (sgRNA) sites (termed sg1-5) that would be expected to cleave one α-globin gene and not the other **(Fig. 1A; Supplemental Fig. 2a & b)**. We therefore chose to test guides that exploited sequence differences between the two genes that targeted the 3’ UTR. To do so, we delivered each chemically-modified sgRNA^33^ pre-complexed with Cas9 ribonucleoprotein (RNP) to human CD34^+^ HSPCs by electroporation in order to determine which of the five 3’ UTR guides can most efficiently and specifically induce indels (insertions/deletions) at the intended gene. We then PCR amplified the 3’ UTR regions of both *HBA2* and *HBA1* and analyzed the corresponding Sanger sequences for indel frequencies using TIDE analysis^34^. We found four of the five guides to induce indels at high frequencies, two of which were capable of distinguishing between the two genes (sg2 cuts *HBA2* and sg5 cuts *HBA1*) and two of which that were not (sg1 and sg4 cut both *HBA2* and *HBA1*)**(Fig. 1B)**. *HBA1* and *HBA2* target sites for sg1 and sg4 only differed by one base pair, likely accounting for the lack of specificity. On the other hand, target sites for the highly-specific sg2 and sg5 differed by five base pairs **(Supplemental Fig. 2a)**. Additionally, to determine the off-target activity of the most effective of these sgRNAs, sg5, we performed targeted sequencing of the forty most likely off-target sites as predicted by COSMID^35^. This determined that the *HBA1*-specific sg5 was extremely specific, with a median on-target activity of 62.9% and only two of the forty predicted sites showing activity above detection threshold across three separate HSPC donors (median of 0.24% and 0.13% activity at off-target sites 1 and 12, respectively)**(Supplemental Fig. 3)**.

**Figure 1:**
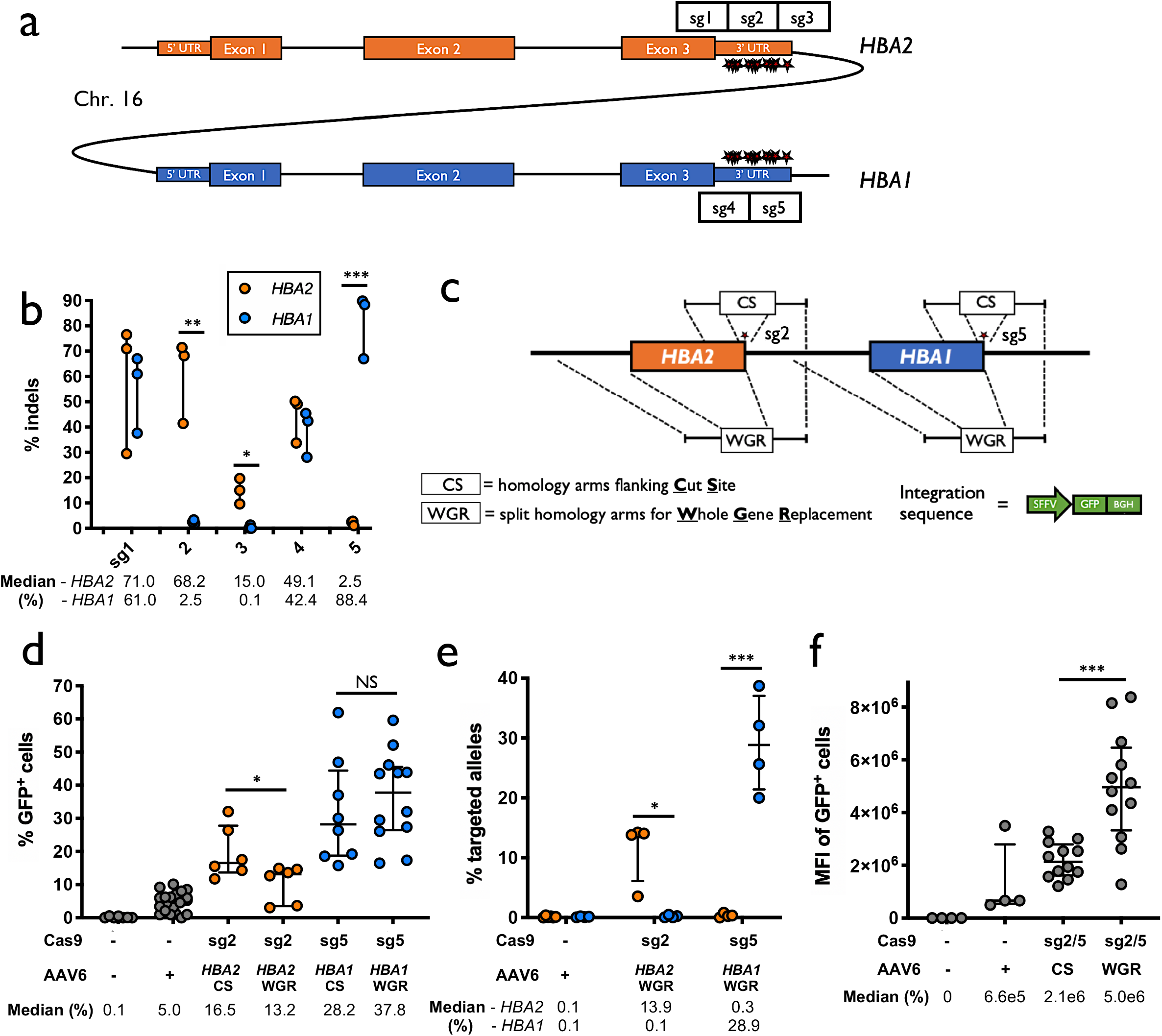
sgRNA & AAV6 design for CRISPR/AAV6-mediated targeting of α-globin locus. A) Schematic of *HBA2* and *HBA1* genomic DNA. Sequence differences between the two genes in 3’ UTR region are depicted as red stars. Locations of the five prospective sgRNAs are indicated. B) Indel frequencies for each guide at both *HBA2* and *HBA1* in human CD34^+^ HSPCs are depicted in orange and blue, respectively. Bars represent median + interquartile range. N = 3 for each treatment group. *: P<0.05; **: P<0.005; ***: P<0.0005 determined using unpaired t test. C) AAV6 DNA repair donor design schematics to introduce a SFFV-GFP-BGH integration are depicted at *HBA2* and *HBA1* loci. D) Percentage of GFP^+^ cells using *HBA2*- and *HBA1*-specific guides and CS and WGR SFFV-GFP AAV6 donors as determined by flow cytometry. Bars represent median + interquartile range. N > 6 for each treatment group. *: P<0.05 determined using unpaired t test. E) Targeted allele frequency at *HBA2* and *HBA1* as determined by ddPCR, to determine whether off-target integration occurs at the unintended gene. Bars represent median + interquartile range. N > 4 for each treatment group. *: P<0.05; ***: P<0.0005 determined using unpaired t test. F) MFI of GFP^+^ cells across each targeting event as determined by BD FACSAria II platform. Bars represent median + interquartile range. N > 4 for each treatment group. ***: P<0.0005 determined using unpaired t test.

### Split homology arm strategy allows whole gene replacement of α-globin

Upon identification of *HBA1-* and *HBA2*-specific guides, we proceeded to test the frequency of AAV6-mediated homologous recombination (HR) at these loci. To do so, we designed AAV6 repair template vectors that would allow us to integrate a GFP expression cassette directly into the sites of the Cas9 RNP-induced breaks. This was facilitated by 400bp homology arms immediately flanking the cut site (hereafter termed “CS”) of each sgRNA **(Fig. 1C)**. As before, we delivered the most specific guides (sg2 to target *HBA2* and sg5 to target *HBA1*) complexed with Cas9 RNP by electroporation into human CD34^+^ HSPCs. Because of previously-reported electroporation-aided transduction^36^, we added each AAV6 vector immediately following electroporation of Cas9 RNP in order to maximize AAV delivery. Several days later, after episomal expression in the “AAV only” controls was reduced, we analyzed targeting frequencies by flow cytometry **(Supplemental Fig. 4a)**. As expected, we found that vectors with homology arms flanking the cut site efficiently integrated at both *HBA2* and *HBA1* in CD34^+^ HSPCs as determined by flow cytometry (median of 16.5% and 28.2% cells were GFP^+^, respectively) **(Fig. 1D)**. However, the predicted cut sites for both guides reside 13bp downstream of the stop codon and the indel spectrum for sg5 indicates that 99.8% of detected indels occur within 9bp of the cut site **(Supplemental Fig. 2c)**. Consequently, this approach would not ensure knockout of either α-globin gene upon HR. Therefore we also cloned repair template vectors with a left homology arm split off from the cut site, spanning the immediate 400bp upstream of each gene. This approach exploits the HR repair process and could facilitate full replacement of the coding region of each α-globin gene, not only reducing α-globin production, but also allowing expression of the transgene to be driven by the endogenous α-globin promoter (hereafter termed “WGR” for “whole gene replacement”)**(Fig. 1C)**. We found that splitting off the left homology arm in the WGR strategy significantly reduced editing frequency at *HBA2* (median of 16.5% vs. 13.2%; P<0.05)**(Fig. 1D)**. Surprisingly, this effect appeared to be gene-dependent, as the WGR strategy at *HBA1* did not yield a coordinate decrease in editing frequency (median of 28.2% vs. 37.8%)**(Fig. 1D)**. Because the left homology arms were identical in each WGR vector, we used droplet digital PCR (ddPCR) to confirm that our editing events were specific only into the intended gene and correlated well with our targeting frequencies as determined by GFP expression **(Fig. 1E)**. Notably, when performing flow cytometry on these cells, we found that the mean fluorescence intensity (MFI) of GFP^+^ cells was significantly higher for WGR vectors compared to CS vectors (P<0.0005)**(Fig. 1F; Supplemental Fig. 4b**).

### Whole gene replacement at α-globin yields RBC-specific transgene expression

Due to the fact that the WGR repair template design at *HBA1* 1) leads to equivalent editing frequencies compared to the CS design, 2) yields GFP expression levels that are greater than the CS design, and 3) ensures the knockout of one gene copy of α-globin, we next adapted this scheme to replace the HBA1 locus with a full-length *HBB* transgene. To facilitate tracking of *HBB* transgene expression we fused it to a T2A-YFP sequence, which enables fluorescent readout of editing frequencies and a surrogate for *HBB* protein levels **(Fig. 2A)**. To determine the significance of untranslated regions (UTRs) flanking the HBB-T2A-YFP cassette (either *HBB* UTRs or endogenous *HBA1/2* UTRs), as well as the impact of removing the largest *HBB* intron (intron 2, 850bp), we designed multiple different AAV6 repair template vectors and analyzed gene replacement frequencies and transgene expression. For the vectors shown, this means that for the two constructs with *HBB* UTRs, the 5’ UTR of *HBA1* would be replaced by that of HBB, while the 3’ UTR of *HBB* would be introduced followed by the 3’ UTR of *HBA1*. For the *HBA*-targeting vectors, the UTRs included in the AAV donor would simply act as longer homology arms and lead to gene replacement with the full-length *HBB* (including introns) from start codon to stop codon—effectively maintaining endogenous UTR regions. We targeted HSPCs as previously described, then differentiated cells into erythrocytes using an established protocol^37,38^ and determined targeting frequencies and expression levels by flow cytometry (**Supplemental Fig. 5 & 6a)**. We found that targeting HSPCs at *HBA1* and *HBA2* had no discernible bearing on their ability to differentiate into RBCs compared to “Mock” (i.e. electroporation only), “RNP only”, and “AAV only” controls **(Fig. 2B)**. Targeting frequencies were confirmed by both flow cytometry and ddPCR, allowing us to conclude that the vector with *HBA1* UTRs most efficiently mediated editing (a median of 55.4% of cells were YFP^+^, and 24.4% of total alleles in the bulk population were targeted)**(Fig. 2C and 2D)**. Furthermore, we found that the MFI of YFP^+^ cells was significantly greater for the *HBA1* UTR vector compared to either vector with *HBB* UTRs (P<0.05)**(Fig. 2E; Supplemental Fig. 6b)**, potentially indicating higher *HBB* expression levels in the context of *HBA1* regulatory regions. Because the HBB-T2A-YFP cassette is driven by the endogenous promoter, we were able to determine that YFP was only expressed in GPA^+^/CD71^+^ RBCs **(Fig. 2F)**, leading us to conclude that *HBA1* is an effective safe harbor site for achieving RBC-specific expression, while leaving α-globin production from *HBA2* unperturbed.

**Figure 2:**
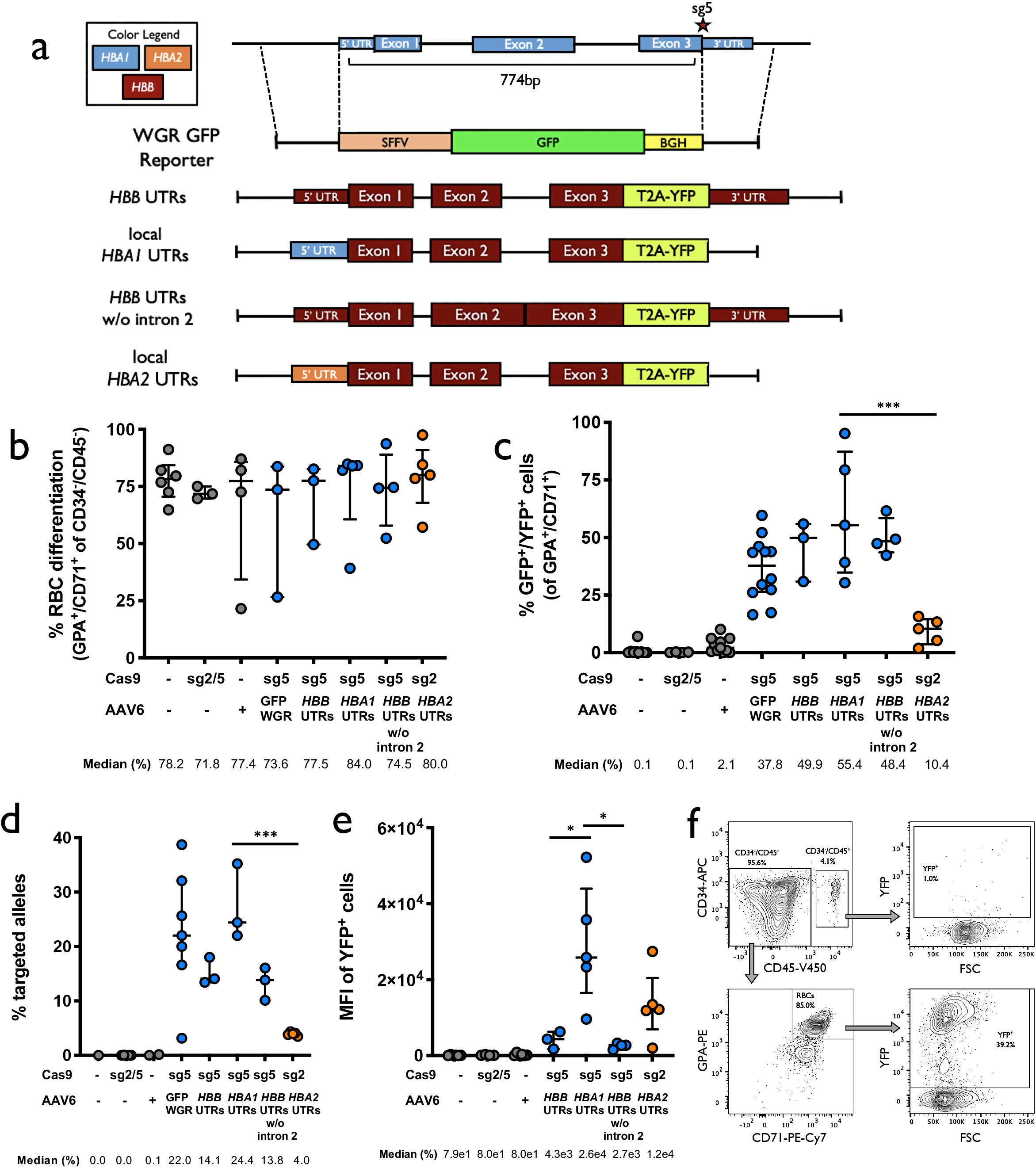
CRISPR/AAV6-mediated targeting of the α-locus using a T2A scheme. A) AAV6 DNA repair donor design schematics to introduce a HBB-T2A-YFP integration are depicted at the *HBA1* locus. B) Percentage of CD34^−^/CD45^−^ HSPCs that acquire RBC surface markers, GPA and CD71, as determined by flow cytometry. Bars represent median + interquartile range. N > 3 for each treatment group. C) Percentage of GFP^+^ cells using *HBA2*- and *HBA1*-specific guides and HBB-T2A-YFP AAV6 donors as determined by flow cytometry. Bars represent median + interquartile range. N > 3 for each treatment group. **: P<0.005 determined using unpaired t test. D) Targeted allele frequency in bulk edited population at *HBA2* and *HBA1* as determined by ddPCR. Bars represent median + interquartile range. N > 3 for each treatment group. ***: P<0.0005 determined using unpaired t test. E) MFI of GFP^+^ cells across each targeting event as determined by BD FACSAria II platform. N > 3 for each treatment group. Bars represent median + interquartile range. F) Representative flow cytometry staining and gating scheme for human HSPCs targeted at *HBA1* with HBB-T2A-YFP (*HBA1* UTRs) and differentiated into RBCs over the course of a 14-day protocol. This indicates that only RBCs (CD34^−^/CD45^−^/CD71^+^/GPA^+^) are able to express the integrated T2A-YFP marker. Analysis was performed on BD FACS Aria II platform.

### Gene replacement of HBA1 with HBB transgene yields adult hemoglobin tetramers

To confirm production of β-globin protein following targeted replacement of *HBA1* with our *HBB* transgene, we screened a number of AAV6 vectors with our *HBB* transgene alone without a T2A-YFP. These vectors used various combinations of regulatory elements, such as *HBB* and *HBA1* 3’ UTRs, WPREs, and BGH poly A regions, as well as a variety of vectors expressing *tNGFR* that would enable us to identify and enrich for a population of highly edited cells **(Fig. 3A; Supplemental Fig. 7)**. We also created gene replacement vectors with lengthened left and right homology arms, hypothesizing that doing so could help the cell identify regions of homology— particularly within the left arm that is split off from the cut site—and thereby increase the editing frequency of our *HBB* transgene. To screen these vectors, we targeted SCD-derived CD34^+^ HSPCs because they exclusively express sickle hemoglobin (HgbS) enabling us to determine degree of adult hemoglobin (HgbA) rescue that results from our editing scheme. HSPCs were then differentiated into RBCs and analyzed for targeting frequency and their ability to form HgbA tetramers using ddPCR and HPLC, respectively. This comparative analysis determined that vectors depicted in **Fig. 3A** were best able replace the *HBA1* locus and yield HgbA tetramers. These experiments indicated that editing at the *HBA1* locus had no significant effect on cell viability 2-4d post-editing **(Supplemental Fig. 8)** or on the or the ability of cells to differentiate into RBCs **(Fig. 3B)**. We also found that gene replacement frequency was significantly improved by lengthening the homology arms, increasing targeted alleles from a median of 21.1% to 36.5% in the bulk population (P<0.05)**(Fig. 3C)**, which is expected to correspond to 48.7% of cells having undergone at least one editing event **(Supplemental Fig. 9)**. When RBCs were analyzed for human hemoglobin by HPLC, we found that all three vectors were able to express and form HgbA tetramers **(Fig. 3D)**. As predicted by the HBB-T2A-YFP vectors, gene replacement cassettes that left local *HBA1* UTRs intact yielded a greater amount of HgbA tetramers, indicating that the T2A-YFP system is highly predictive of transgene expression. We also found that the vector with elongated homology arms not only resulted in significantly higher editing frequencies than the vector with 400bp homology arms, but yielded a significantly greater percentage of HgbA tetramers as well (P<0.05)**(Fig. 3E)**. Importantly, because the vector with the elongated homology arms introduces an identical genome editing event to the *HBA1* UTR vector with 400bp homology arms, we expect this increase in HgbA production to simply be due to the greater frequency at which the long homology arm vector is able to target this locus. In fact, we found a strong correlation between targeting frequency and HgbA tetramer production (R^2^ = 0.8695)**(Fig. 3F)**, indicating that every *HBB*-targeted *HBA1* allele is contributing to endogenous protein levels.

**Figure 3:**
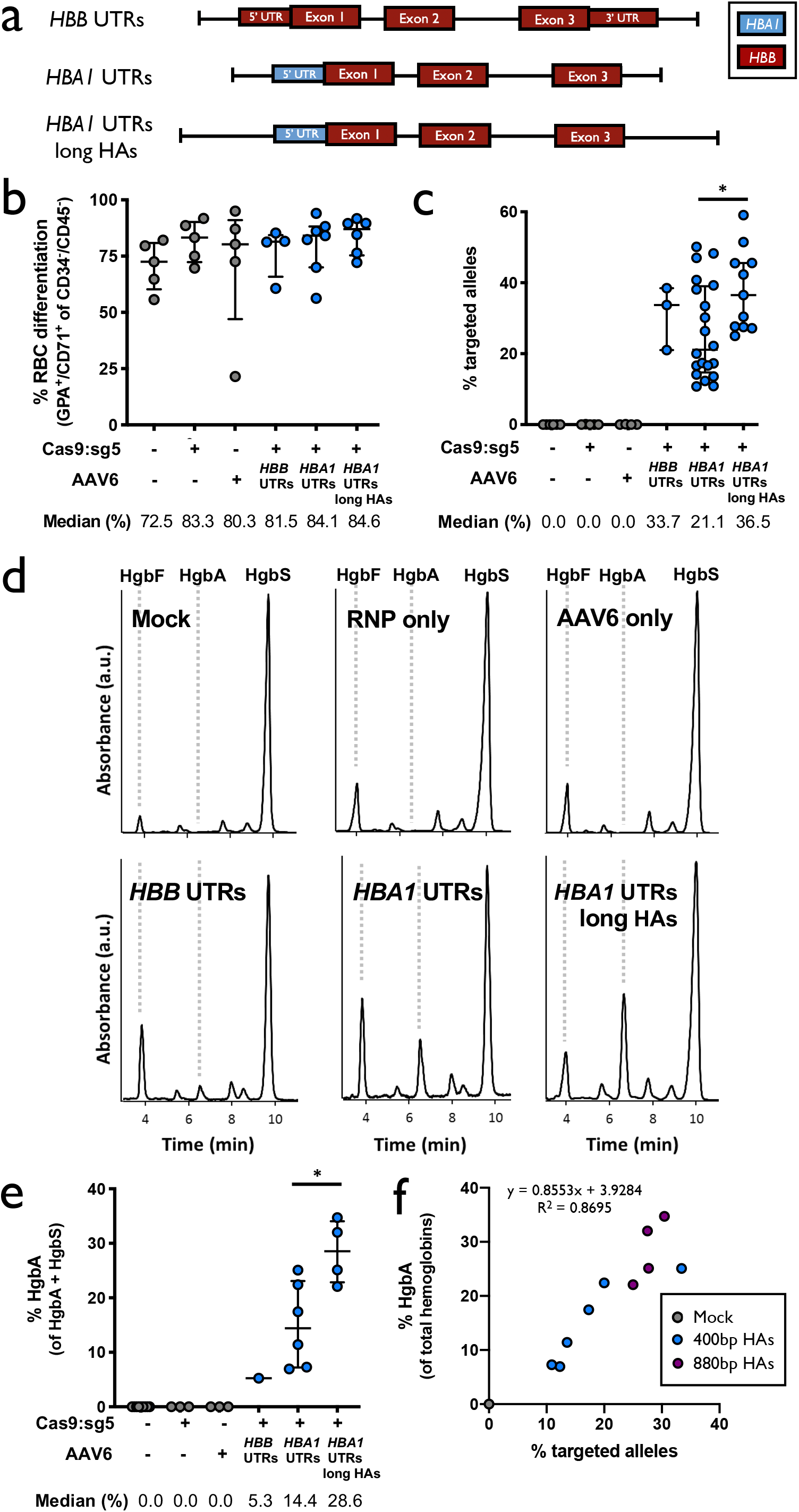
CRISPR/AAV6-mediated targeting at the α-globin locus in SCD HSPCs. A) AAV6 DNA repair donor design schematics to introduce a whole gene replacement *HBB* transgene integration at the *HBA1* locus. B) Percentage of CD34^−^/CD45^−^ HSPCs that acquire RBC surface markers, GPA and CD71, as determined by flow cytometry. Bars represent median + interquartile range. N > 4 for each treatment group. C) Targeted allele frequency in bulk edited population at *HBA1* as determined by ddPCR. Bars represent median + interquartile range. N > 3 for each treatment group. *: P<0.05 determined using unpaired t test. D) Representative HPLC plots for each treatment following targeting and RBC differentiation of human SCD CD34^+^ HSPCs. Retention time for HgbF, HgbA, and HgbS tetramer peaks are indicated. E) Summary of all HPLC results showing percentage of HgbA out of total hemoglobin tetramers. Bars represent median + interquartile range. N > 3 for each treatment group with exception of *HBB* UTRs AAV6 with N = 1. *: P<0.05 determined using unpaired t test. F) Plot depicting correlation between % HgbA vs. % targeted alleles in *HBA1* UTR-targeted samples that were differentiated into RBCs and analyzed by HPLC. Colors of respective vectors are in depicted in figure legend. R^2^ value and trendline formula are indicated. N = 11.

### HSPCs targeted with HBB at HBA1 are capable of long-term engraftment and hematopoietic reconstitution in NSG mice

To determine whether the editing process impacts the ability of HSPCs to engraft and reconstitute myeloid and lymphoid lineages *in vivo*, we performed bone marrow transplantation experiments of human HSPCs targeted at the *HBA1* locus into immune-compromised NSG mice. In order to replicate the clinical HSCT process as closely as possible, HSPCs from healthy donors were mobilized using G-CSF and Plerixafor^12^. Mobilized peripheral blood was collected and HSPCs were then enriched using the CD34 marker and targeted at *HBA1* as above. Two days posttargeting, live CD34^+^ HSPCs were single-cell-sorted into 96-well plates containing methylcellulose media and scored for colony formation ability after incubation for 14d. This indicated that edited HSPCs were able to give rise to cells of all lineages **(Supplemental Fig. 10a-c)**. Although the editing process appears to reduce the total number of colonies, the reduction is primarily due to ability to form colonies in the granulocyte/macrophage lineage, without impacting multi-lineage and erythroid lineage colony formation and their relative distribution.

The entire bulk population of edited cells not used toward the colony-forming assay were injected intrafemorally into immunodeficient NSG mice that had been irradiated in order to clear the hematopoietic stem cell niche in the bone marrow **(Supplemental Fig. 11)**. The experiment was performed on three separate healthy HSPC donors, and due to variable expansion rates among these, the total numbers of cells for transplantation varied among replicates. We therefore designated these as large, medium, and small doses, corresponding to an injection of 1.2 million, 750,000, and 250,000 cells injected per mouse, respectively.

Sixteen weeks post-transplantation, bone marrow from these mice was harvested and engraftment of human cells was determined using human HLA-A/B/C as a marker **(Supplemental Fig. 12)**. We found that all three dosages among all treatment groups were able to successfully engraft into the bone marrow **(Fig. 4A)**. We found that among the medium and small doses, engraftment ability of cells was negatively impacted in AAV only and RNP+AAV treatments compared to Mock electroporated and RNP only controls, as observed previously^21,24^. However, when a greater number of cells were transplanted, we no longer observed any significant differences in engraftment among the treatment groups. We also found that the editing process did not affect the ability of human HSPCs to reconstitute myeloid and lymphoid lineages *in vivo*, and had no discernible impact on the distribution within these lineages among engrafted human cells **(Fig. 4B)**. We next used ddPCR to determine the frequency of the desired targeting event within the population of cells that engrafted. We found that a mean of 11.0% of total alleles within our bulk population of successfully-engrafted HSPCs were properly targeted **(Fig. 4C),** which would be expected to correspond to 18.2% of cells having undergone at least one editing event **(Supplemental Fig. 9)**. We also lineage-sorted engrafted cells into CD19^+^ (B-Cell), CD33^+^ (myeloid), and Lin^−^/CD34^+^/CD10^−^ (HSPC) populations and determined targeting frequencies among these established subpopulations, which was 7.8%, 14.9%, and 17.2%, respectively. We observed a modest reduction in targeted allele frequency from the *in vitro*, pre-transplantation population to the cell population that successfully engrafted **(Fig. 4D)**, which is in line with that observed in previous reports ^22,24^ and much less severe of a drop than recently reported by Pattabhi, *et al*. ^25^. In addition to editing with the clinically-relevant *HBB* gene replacement vector, we also targeted cells with a WGR repair template cassette to replace the *HBA1* gene with a GFP expressed by the strong UbC promoter **(Supplemental Fig. 13a-e)**. We found engraftment rates of human cells at a median 8.7%, indicating that replacement of the *HBA1* gene has little effect on the ability of HSPCs to engraft. We also used flow cytometry to determine editing frequencies of successfully-engrafted cells to be a median of 25.6%, and in CD19^+^, CD33^+^, and Lin^−^ /CD34^+^/CD10^−^ lineages at a mean of 1.0%, 15.9%, and 0.9%, respectively, indicating that edited cells were capable of engraftment and reconstitution of the various lineages.

**Figure 4:**
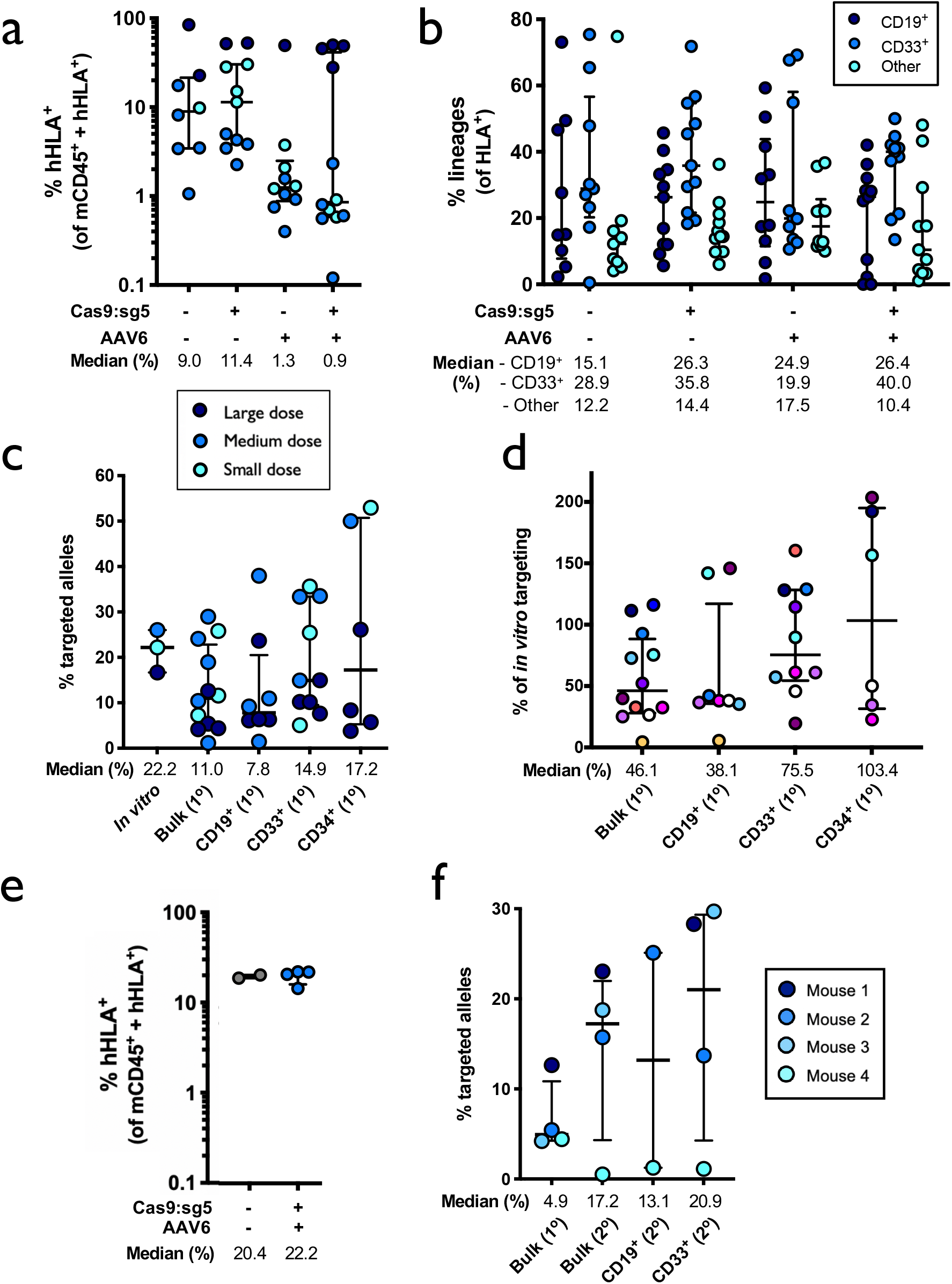
Engraftment of α-globin-targeted human HSPCs into NSG mice. A) 16 weeks after bone marrow transplantation of targeted human CD34^+^ HSPCs into NSG mice, bone marrow was harvested and rates of engraftment were determined. Depicted is percentage of mTerr119^−^ cells (non-RBCs) that were hHLA^+^ from the total number of cells that were either mCd45^+^ of hHLA^+^. Indicated by color coding are large, medium, and small dose experiments where 1.2M, 750K, or 250K cells were initially transplanted, respectively. Bars represent median + interquartile range. N > 8 for each treatment group. B) Among engrafted human cells, the distribution among CD19^+^ (B-cell), CD33^+^ (myeloid), or other (i.e. HSPC/RBC/T/NK/Pre-B) lineages are indicated. Bars represent median + interquartile range. N > 8 for each treatment group. C) Targeted allele frequency at *HBA1* as determined by ddPCR among *in vitro* (pretransplantation) targeted HSPCs and bulk engrafted HSPCs as well as among CD19^+^ (B-cell), CD33^+^ (myeloid), and CD34^+^ (HSC) lineages. Bars represent median + interquartile range. N = 3 for *in vitro* HSPCs, N = 12 for bulk engrafted HSPCs, N = 8 for CD19^+^ HSPCs, N = 10 for CD33^+^ HSPCs, and N = 6 for CD34^+^ HSPCs. D) Targeted allele frequency at *HBA1* among engrafted human cells compared to the bulk targeting rate of the pre-transplantation, *in vitro* human HSPC population. Each mouse is represented by a different color. Bars represent median + interquartile range. N = 12 for bulk engrafted HSPCs, N = 8 for CD19^+^ HSPCs, N = 10 for CD33^+^ HSPCs, and N = 6 for CD34^+^ HSPCs. E) Following primary engraftments, engrafted human cells were transplanted a second time into the bone marrow of NSG mice. 16 weeks post-transplantation, bone marrow was harvested and rates of of engraftment were determined. Depicted is the percentage of mTerr119^−^ cells (non-RBCs) that were hHLA^+^ from the total number of cells that were either mCd45^+^ or hHLA^+^. Bars represent median + interquartile range. N = 2 for mock treatment group and N = 4 for targeted treatment groups. F) Targeted allele frequency at *HBA1* as determined by ddPCR among engrafted human cells in bulk sample as well as among CD19^+^ (B-cell) and CD33^+^ (myeloid) lineages in secondary transplantation experiments. Each mouse is represented by a different color. Bars represent median + interquartile range. N = 4 for each treatment group with exception of CD19^+^ HSPCs with N = 2.

After harvesting the cells that successfully engrafted in the initial transplantation experiment, we injected these cells intravenously into new mice for a second round of engraftment to determine whether the editing process impacts the ability of cells to engraft and repopulate the hematopoietic system long term in a secondary mouse. Indeed, both the control Mock electroporation treatment as well as cells targeted with Cas9 RNP+AAV6 were able to engraft at >20% **(Fig. 4E)**. We then used ddPCR as before to determine the gene replacement frequency within the population of human cells that were able to successfully engraft in this second round of transplantation. In doing so, we observed editing rates in the bulk sample and within the lineages that were in line with the rates observed among cells that engrafted in the initial transplantation experiment **(Fig. 4F)**. These results were further confirmed by targeting with our WGR GFP vector, which also demonstrated that edited (GFP^+^) cells were able to engraft long term when bone marrow was harvested from secondary mice **(Supplemental Fig. 13f-g)**.

### Gene replacement with HBB transgene in ϐ-thalassemia-derived HSPCs corrects ϐ-globin:β-globin imbalance

After demonstrating our ability to achieve stable gene replacement frequencies at the *HBA1* locus in long-term repopulating HSCs derived from WT donors, we sought to determine the effects of our methodology on β-thalassemia-derived HSPCs. To do so, CD34^+^ cells were isolated from back-up Plerixafor- and Filgrastim-mobilized peripheral blood saved from β-thalassemia patients undergoing allogeneic HSCT. As previously, we expanded and targeted these HSPCs using the *HBA1* UTR vectors **(Fig. 3A)** and sorted single live CD34^+^ HSPCs into each well of 96-well plates for colony formation assays. As before, we found that edited HSPCs were able to give rise to cells of all lineages **(Supplemental Fig. 10d & e)**. Although no lineage skewing was apparent, the overall ability of edited β-thalassemia-derived HSPCs to form colonies appeared to be slightly reduced, which is in line with prior reports following Cas9/AAV6-mediated genome editing^39^.

In addition to colony formation assays, a subset of targeted HSPCs were subjected cells to RBC differentiation 2d post-editing. As before, we found that our approach had no discernible bearing on the cell viability post-editing **(Supplemental Fig. 8)**, or on the ability of β-thalassemia-derived HSPCs to differentiate into RBCs (Fig. 5A). As shown previously, we also found that elongating the homology arms significantly improved editing frequencies in these cells as determined by ddPCR (13.8% vs. 45.5%; P<0.005)**(Fig. 5B)**, which is expected to correspond to 63.8% of cells having undergone at least one editing event **(Supplemental Fig. 9)**. To gain insight into the effect that our editing scheme has on the expression of α- and β-globin, we designed ddPCR primer/probes that allowed us to assess mRNA expression of α-globin (not distinguishing between *HBA1* and *HBA2*) as well as mRNA expression from our *HBB* transgene. When expression was normalized to the RBC markers *GPA*, we found that cells edited with the longer homology arm displayed a decrease in α-globin expression and a significant increase in *HBB* transgene expression compared to the RNP only treatment (P<0.05)**(Fig. 5C; Supplemental Fig. 14a)**.

**Figure 5:**
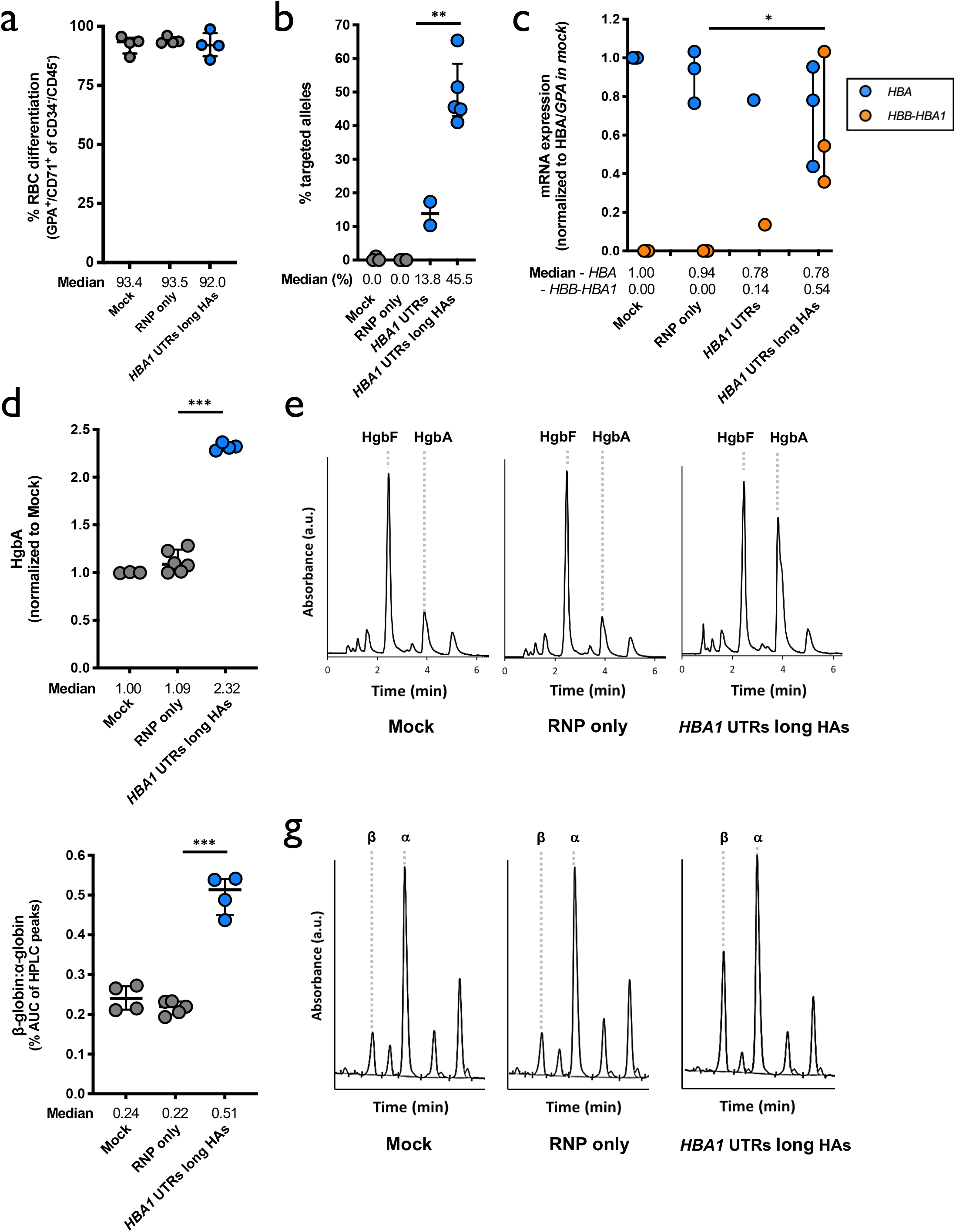
Targeting the α-globin locus in β-thalassemia-derived HSPCs. A) Percentage of CD34^−^/CD45^−^ HSPCs that acquire RBC surface markers, GPA and CD71, as determined by flow cytometry. Bars represent median + interquartile range. N = 4 for each treatment group. B) Targeted allele frequency at *HBA1* in β-thalassemia-derived HSPCs as determined by ddPCR. Bars represent median + interquartile range. N = 3 for mock, N = 2 for RNP only and *HBA1* UTRs, and N = 5 for *HBA1* UTRs long HAs treatments. **: P<0.005 determined using unpaired t test. C) Following differentiation of targeted HSPCs into RBCs, mRNA was harvested and converted into cDNA. Expression of HBA (does not distinguish between *HBA1* and *HBA2*) and *HBB* transgene were normalized to *HBG* expression. Bars represent median + interquartile range. N = 3 for each treatment group with exception of *HBA1* UTRs with N = 1. **: P<0.05 determined using unpaired t test D) Summary of hemoglobin tetramer HPLC results showing HgbA normalized to HgbF. Bars represent median + interquartile range. N > 3 for each treatment group. ***: P<0.0001 determined using unpaired t test. E) Representative hemoglobin tetramer HPLC plots for each treatment following targeting and RBC differentiation of HSPCs. Retention time for HgbF and HgbA tetramer peaks are indicated. F) Summary of reverse-phase globin chain HPLC results showing area under the curve (AUC) of β-globin/AUC of α-globin. Bars represent median + interquartile range. N > 4 for each treatment group. ***: P<0.0001 determined using unpaired t test. G) Representative reverse-phase globin chain HPLC plots for each treatment following targeting and RBC differentiation of HSPCs. Retention time for HgbF and HgbA tetramer peaks are indicated.

Though we do see correction of the extreme globin-chain imbalance at the mRNA level following editing, a more appropriate clinical readout would be whether our correction scheme allows patient-derived RBCs to produce more HgbA tetramers. Therefore, as before, we harvested RBC pellets at d14 of differentiation and performed HgbA tetramer analysis by HPLC. Indeed, we found that cells targeted with our optimized vector with longer homology arms was able to significantly boost HgbA tetramers *in vitro* (P<0.0001)**(Fig. 5D & E)**. In order to determine the effect of our editing strategy at an even greater resolution, we also performed reverse-phase HPLC which allowed us to determine relative β-globin and α-globin levels within our treatment groups. As expected based on the mRNA analysis, we observed a significant correction of the ratio of β-globin to α-globin in our edited cell population (P<0.00001)**(Fig. 5F & G)**. This normalization was attributed to a significant increase in β-globin production (P<0.0001) as well as a modest, but also significant decrease in α-globin (P<0.005)**(Supplemental Fig. 14b)**.

### Targeted β-thalassemia-derived HSPCs are capable of long-term engraftment and hematopoietic reconstitution in NSG mice

In addition to *in vitro* RBC differentiation and analysis, we performed engraftment experiments as before by injecting targeted β-thalassemia-derived HSPCs into irradiated NSG mice. Twelve to sixteen weeks post-transplantation, we harvested bone marrow from mice and determined engraftment and targeting frequencies by flow cytometry and ddPCR, respectively. We found that indeed, patient-derived HSPCs targeted with our transgene at the *HBA1* locus were capable of engraftment, with a median of 3.4% human cells in the bone marrow **(Fig. 6A)**, as well as multi-lineage reconstitution **(Fig. 6B)**. Using ddPCR, we also determined that successfully-engrafted cells were edited in the bulk population at a median frequency of 5.1% as well as in CD19^+^, CD33^+^, and “Other” lineages at median frequencies of 1.4%, 1.4%, and 2.5%, respectively **(Fig. 5C)**.

**Figure 6:**
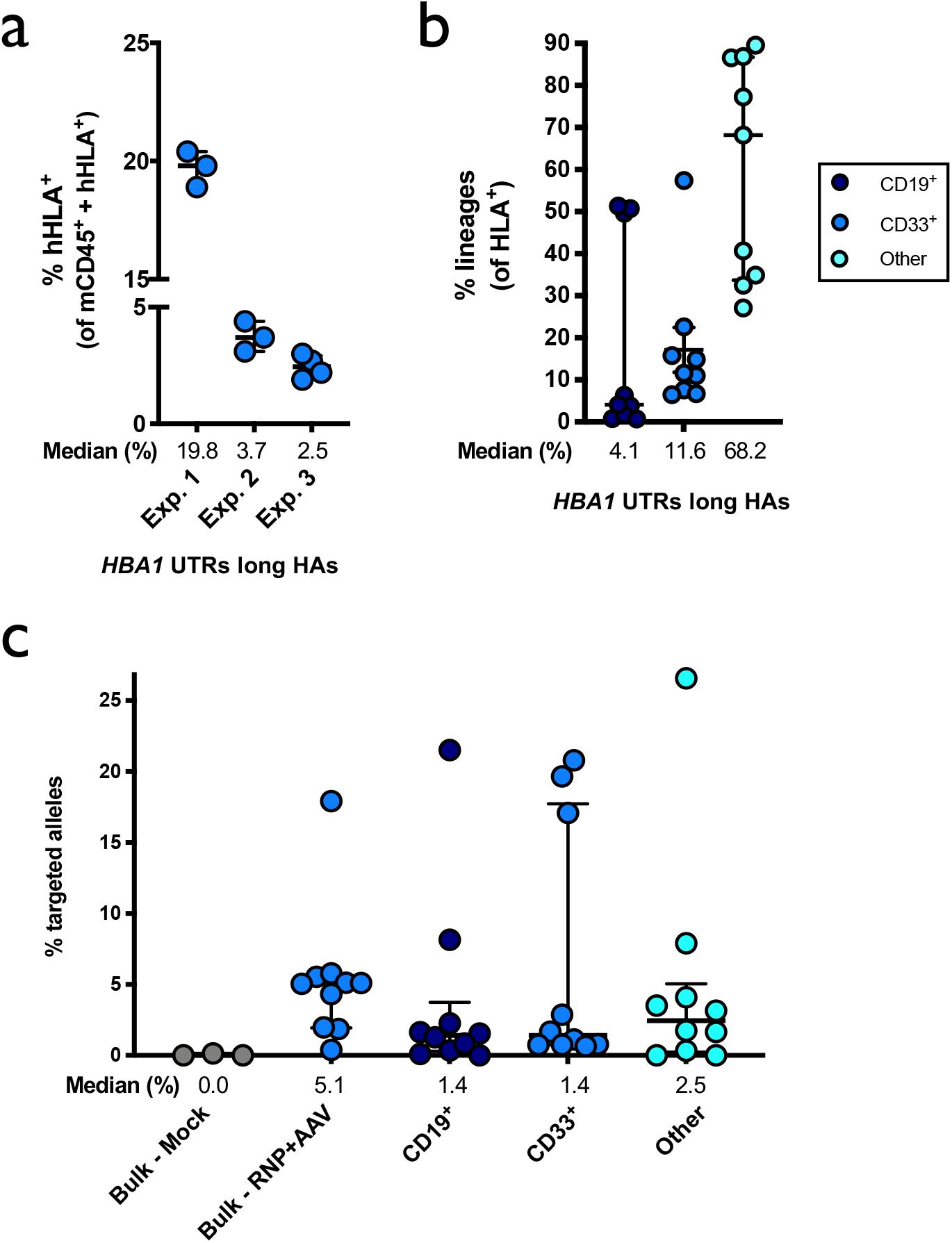
Engraftment of α-globin-targeted β-thalassemia-derived HSPCs into NSG mice. A) 16 weeks after bone marrow transplantation of targeted β-thalassemia-derived HSPCs into NSG mice, bone marrow was harvested and rates of engraftment were determined. Depicted is percentage of mTerr119^−^ cells (non-RBCs) that were hHLA^+^ from the total number of cells that were either mCd45^+^ of hHLA^+^. Bars represent median + interquartile range. N = 10. B) Among engrafted human cells, the distribution among B-cell, myeloid, or other (i.e. HSPC/RBC/T/NK/Pre-B) lineages are indicated. Bars represent median + interquartile range. N = 9. C) Targeted allele frequency at *HBA1* as determined by ddPCR among engrafted human cells in bulk sample as well as among CD19^+^ (B-cell), CD33^+^ (myeloid), and other (i.e. HSPC/RBC/T/NK/Pre-B) lineages in secondary transplantation experiments. Bars represent median + interquartile range. N = 3 for mock treatment group and N = 10 for targeted treatment group.

## DISCUSSION

In summary, the above results have described a novel genome editing protocol for the treatment of β-thalassemia which address both molecular factors responsible for the disease— loss of β-globin and accumulation of excess α-globin—in a single genome editing event. Prior data indicates that approximately 25% edited cell chimerism in the bone marrow appears to be the threshold by which transfusion-independence is achieved in thalassemia patients^40^. Therefore, the editing rates we are achieving in β-thalassemia-derived HSPCs (45.5% of alleles targeted *in vitro*) may be sufficient for correction of the disease. However, it must be noted that we observed a substantial reduction in the editing frequencies of cells that successfully engrafted into NSG mice *in vivo* (5.1% of alleles targeted within this population). While this is consistent with previously reported *ex vivo* genome-editing strategies^21–24^, the apparent reduction in the ability of edited HSCs to undergo editing and/or long-term engraftment currently stands as a barrier to clinical translation. Currently there is much ongoing effort within the field to address this issue, including optimization of cell cycling conditions, small molecules that improve cell viability, and strategies to bias cells from non-homologous end joining toward HDR.

Nevertheless, because our approach is site-specific and uses a patient’s *own* cells, it would: 1) Overcome the shortage of immunologically-matched donors; 2) Eliminate the need for constant blood transfusion and/or iron chelation therapy; 3) Dramatically reduce the likelihood of immune rejection that accompanies allogeneic HSCT; and 4) Avoid the risks of semi-random integration of viral vectors in the genome. For these reasons, the technology we have described helps overcome the pitfalls of the current therapeutic strategies, leading us to conclude that our strategy has the potential to become the new standard of care. However, much work remains to be done in order to translate this work to the clinic. Nevertheless, we believe that the reported results establish an important proof-of-concept for the field of genome-editing as well as for the treatment of hemoglobinopathies.

Beyond the immediate impact for treatment of β-thalassemia, we also believe that our results have broader relevance to the genome editing field as a whole. To our knowledge, the above results represent the first time that a single Cas9-induced DSB was able to facilitate replacement of a large genomic region with a custom editing cassette at high frequencies. Because most recessive hereditary diseases are caused by loss-of-function mutations throughout a particular gene, the scheme that we have developed can be adapted into a one-size-fits-all treatment strategy for a wide range of genetic disorders, effectively expanding the genome editing toolbox.

Our study also showed that the T2A cleavage peptide system coupled with a fluorescent reporter was highly predictive of transgene expression. This demonstrates the utility of this system in rapidly identifying successfully-edited cells and for comparison of a variety of editing vectors (i.e. those with different regulatory regions, with or without specific introns, etc.). Because patient-derived HSPCs are so hard to obtain, especially from multiple donors, this T2A screening system also allows for identification of the optimal translational vector in healthy-HSPCs that can be validated in patient-derived HSPCs. Lastly, because we have optimized gene replacement at the α-globin locus, which is only expressed in RBCs, this work has characterized a safe harbor locus for delivery of payloads by RBCs, such as therapeutic enzymes and monoclonal antibodies. This will allow future work to replace the *HBA1* locus with custom vectors at high frequencies, thereby achieving RBC-specific expression without the risk of knocking out a gene that is critical to RBC development (because *HBA2* remains intact). For these reasons, we expect the findings of this study to guide future genome editing work, both as strategies for correction of a wide range of genetic disorders as well as for a variety of cell engineering applications.

---

**Supplemental Figure 1:**
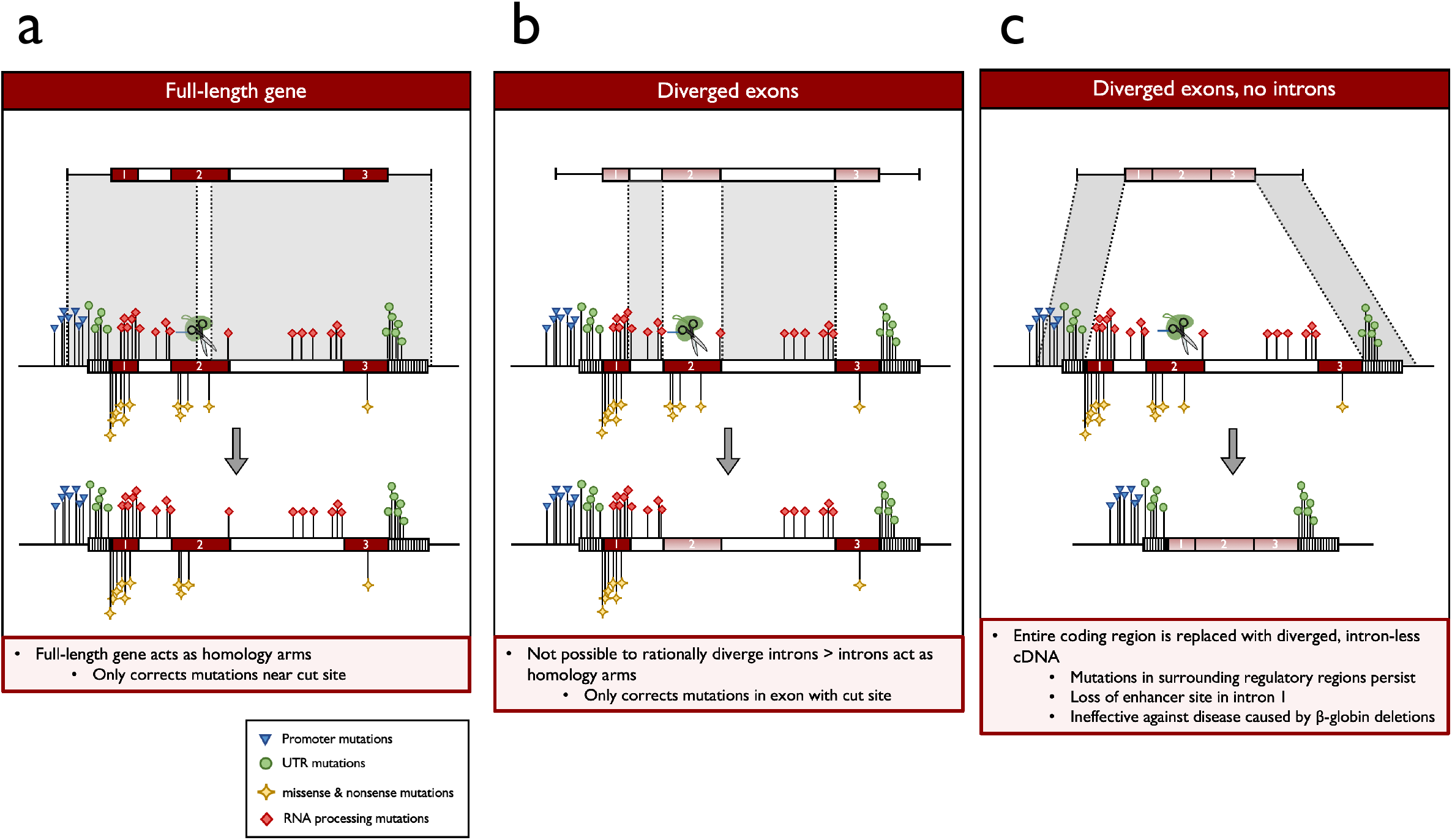
Expected outcomes of introducing *HBB* transgene at endogenous locus. A) Expected outcome when integrating an undiverged, full-length *HBB* (with introns) at the endogenous locus of HSPCs derived from patients with β-thalassemia. The varieties of disease-causing mutations are annotated in the figure legend. B) Expected outcome when integrating a diverged, full-length *HBB* (with introns) at the endogenous locus of HSPCs derived from patients with β-thalassemia. C) Expected outcome when integrating a diverged, *HBB* cDNA (without introns) at the endogenous locus of HSPCs derived from patients with β-thalassemia.

**Supplemental Figure 2:**
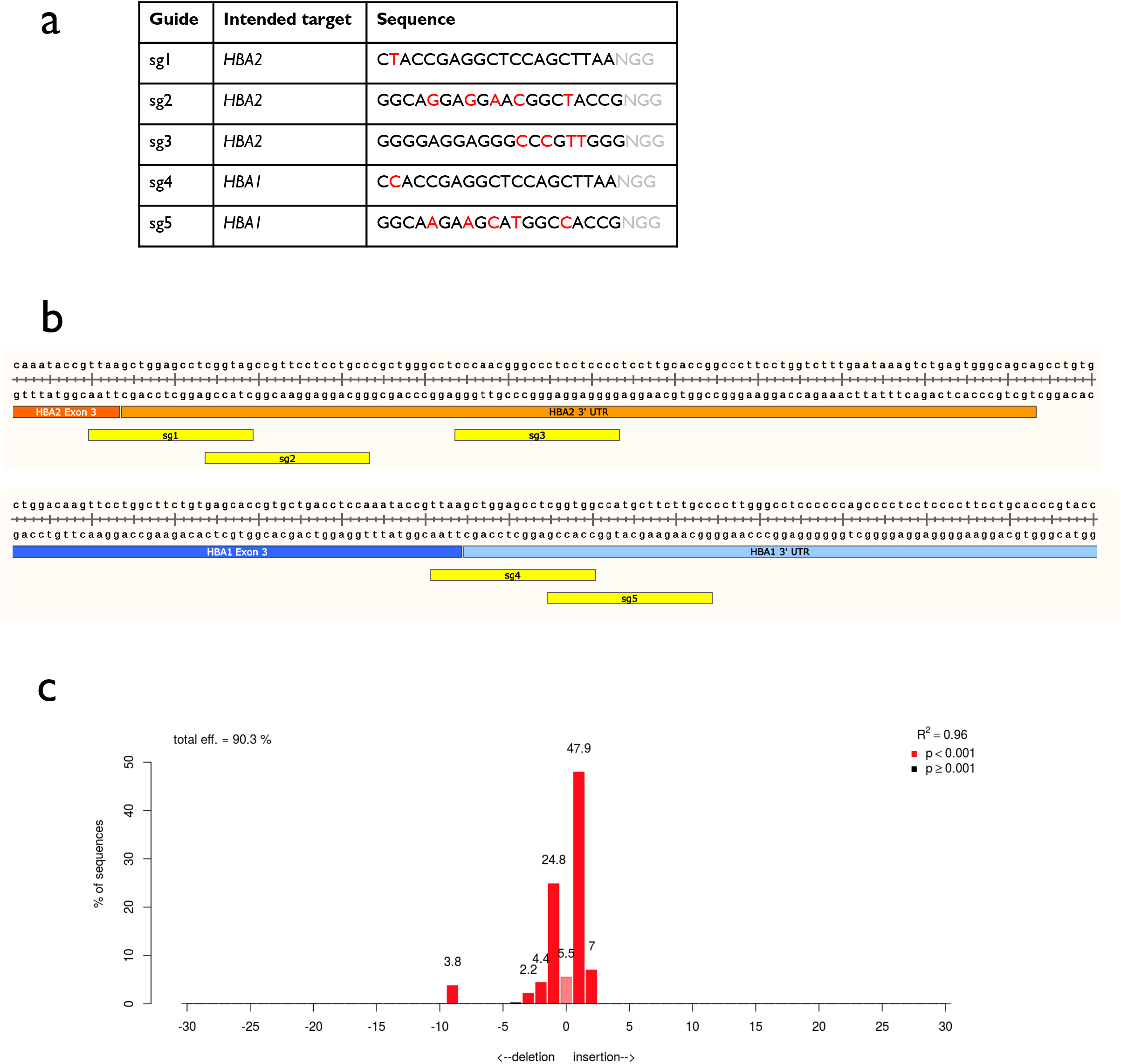
Analysis of Cas9 sgRNAs targeting α-globin loci. A) Table with guide RNA sequences. PAM shown in grey, and differences between *HBA1* and *HBA2* are highlighted in red for each guide. B) Schematic depicting locations of all five guide sequences at genomic loci. C) Representative indel spectrum of *HBA1*-specific sg5 generated by TIDE software.

**Supplemental Figure 3:**
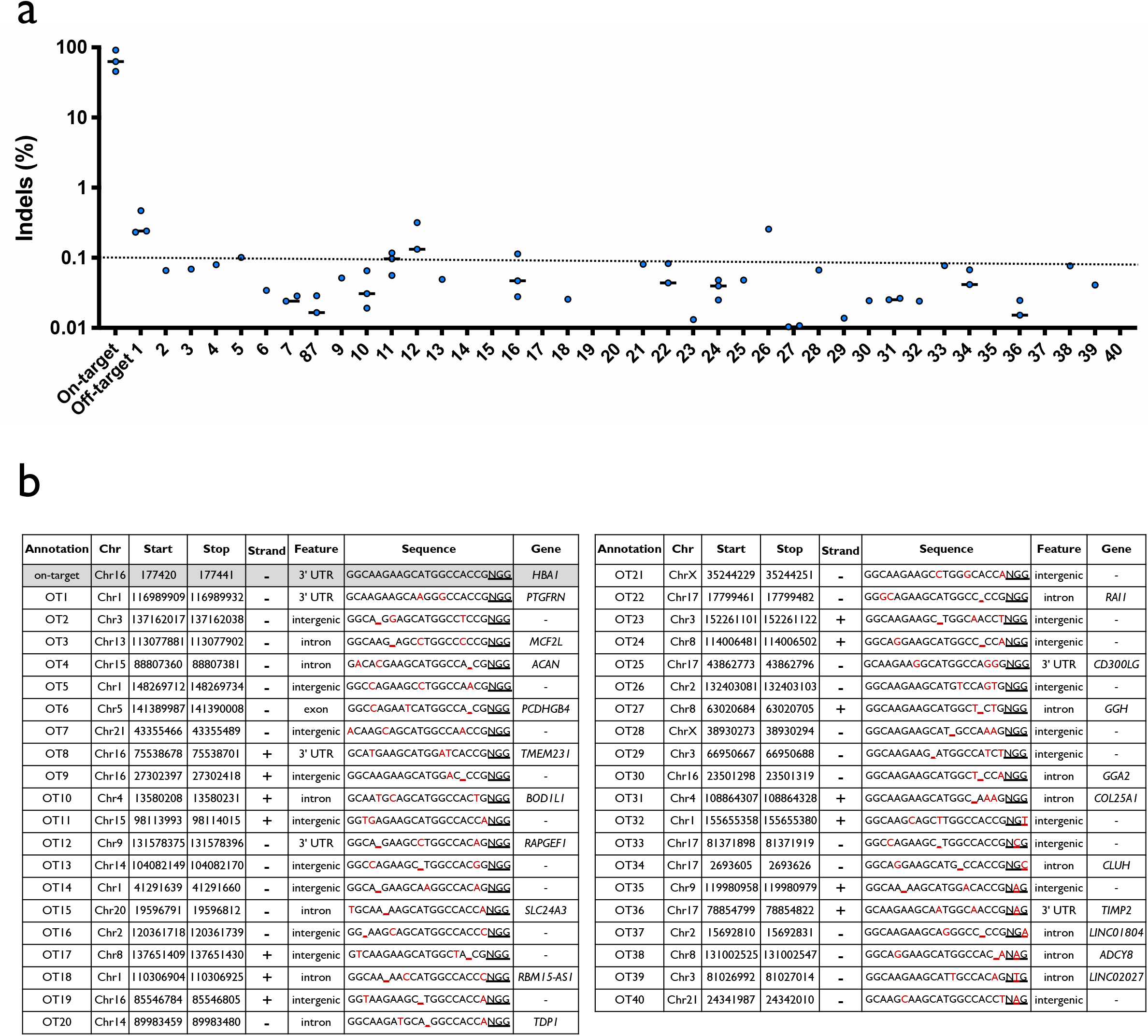
Off-target analysis of *HBA1*-specific sg5. A) Summary of rhAmpSeq targeted sequencing results at on-target and 40 most highly-predicted off-target sites by COSMID for *HBA1* sg5. Values are indel frequency for RNP treatment after subtraction of indel frequency for Mock treatment at each locus for each experimental replicate. N = 3, though not all values are displayed since some were <0.01% after subtraction of Mock indel frequencies. Bars represent median. B) List of genomic coordinates for forty most highly-predicted off-target sites by COSMID for *HBA1* sg5.

**Supplemental Figure 4:**
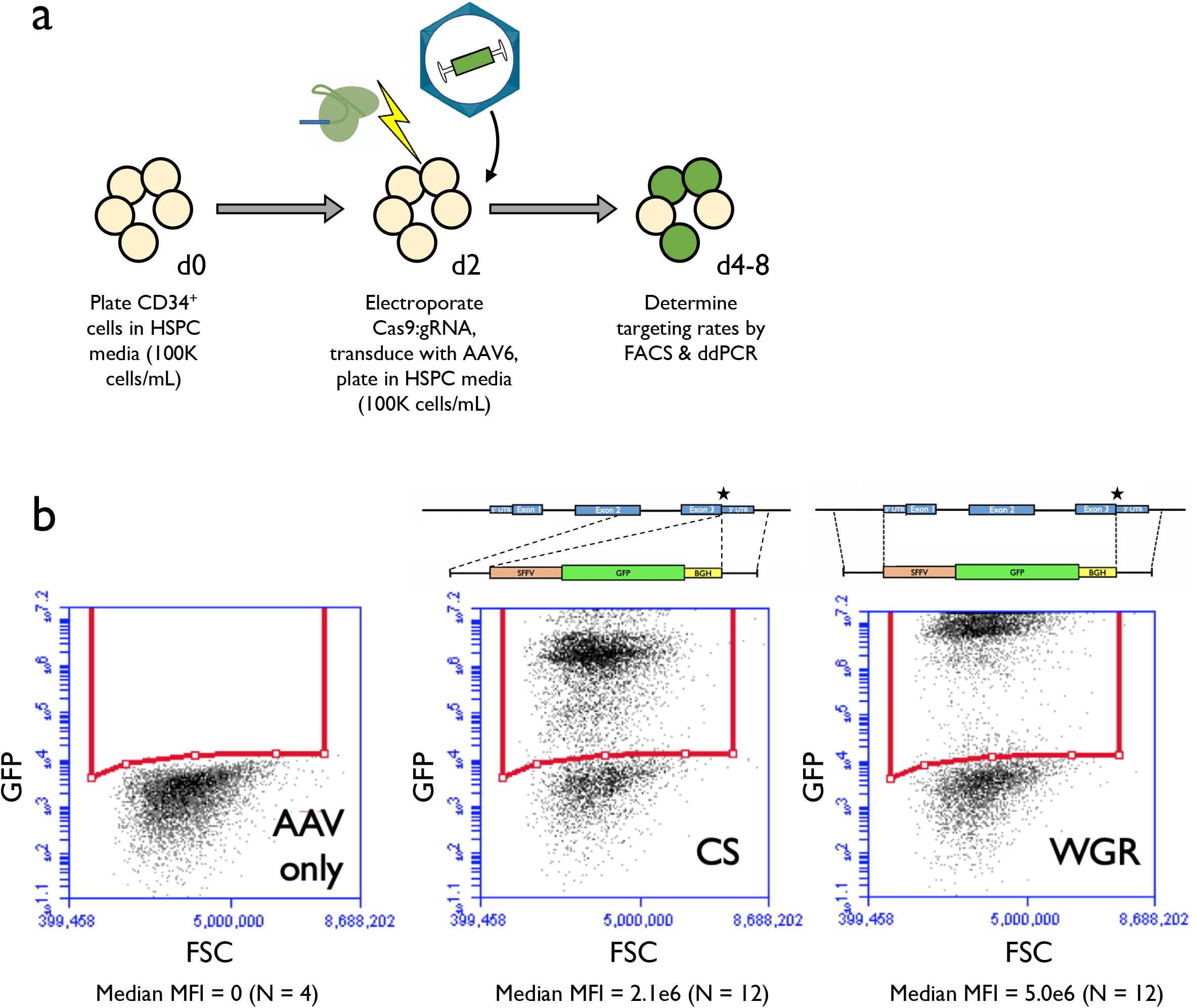
Targeting HSPCs with GFP-*HBA* integration vectors. A) Timeline for editing and analysis of HSPCs targeted with *GFP-HBA* integration vectors. B) Depicted are representative flow cytometry images for human HSPCs that have been targeted by CRISPR/AAV6 methodology 14d post-editing. This indicates that whole-gene-replacement (WGR) integration yields a greater MFI per GFP^+^ cell than cut-site (CS) integration at the *HBA1* locus. Analysis was performed on BD Accuri C6 platform. Median MFI across all replicates is shown below each flow cytometry image, and schematics of integration vectors are shown above.

**Supplemental Figure 5:**
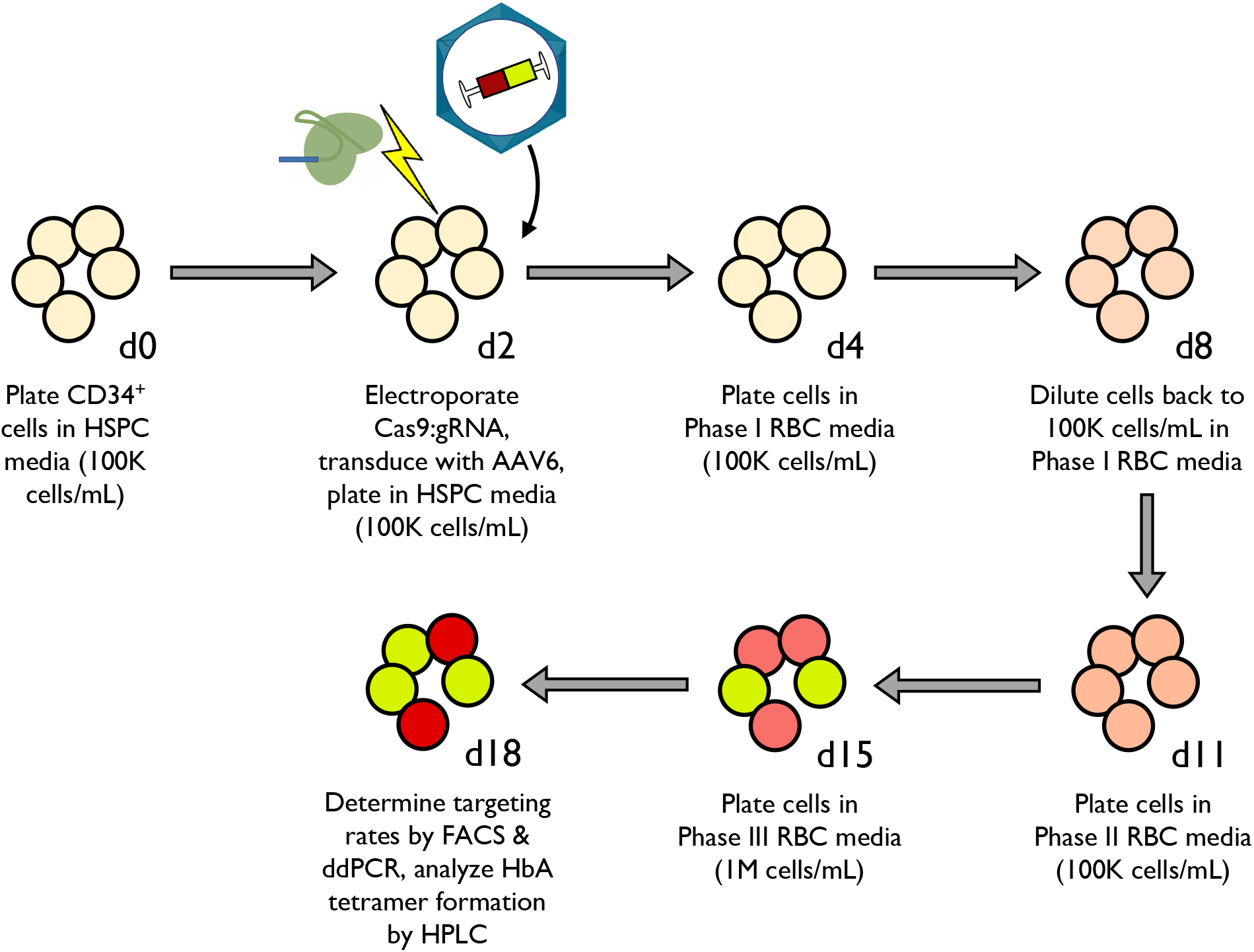
Timeline for targeting HSPCs with HBB-T2A-YFP-*HBA* integration vectors. Timeline for targeting of HSPCs with HBB-T2A-YFP integration vectors, differentiation into RBCs, and subsequent analysis.

**Supplemental Figure 6:**
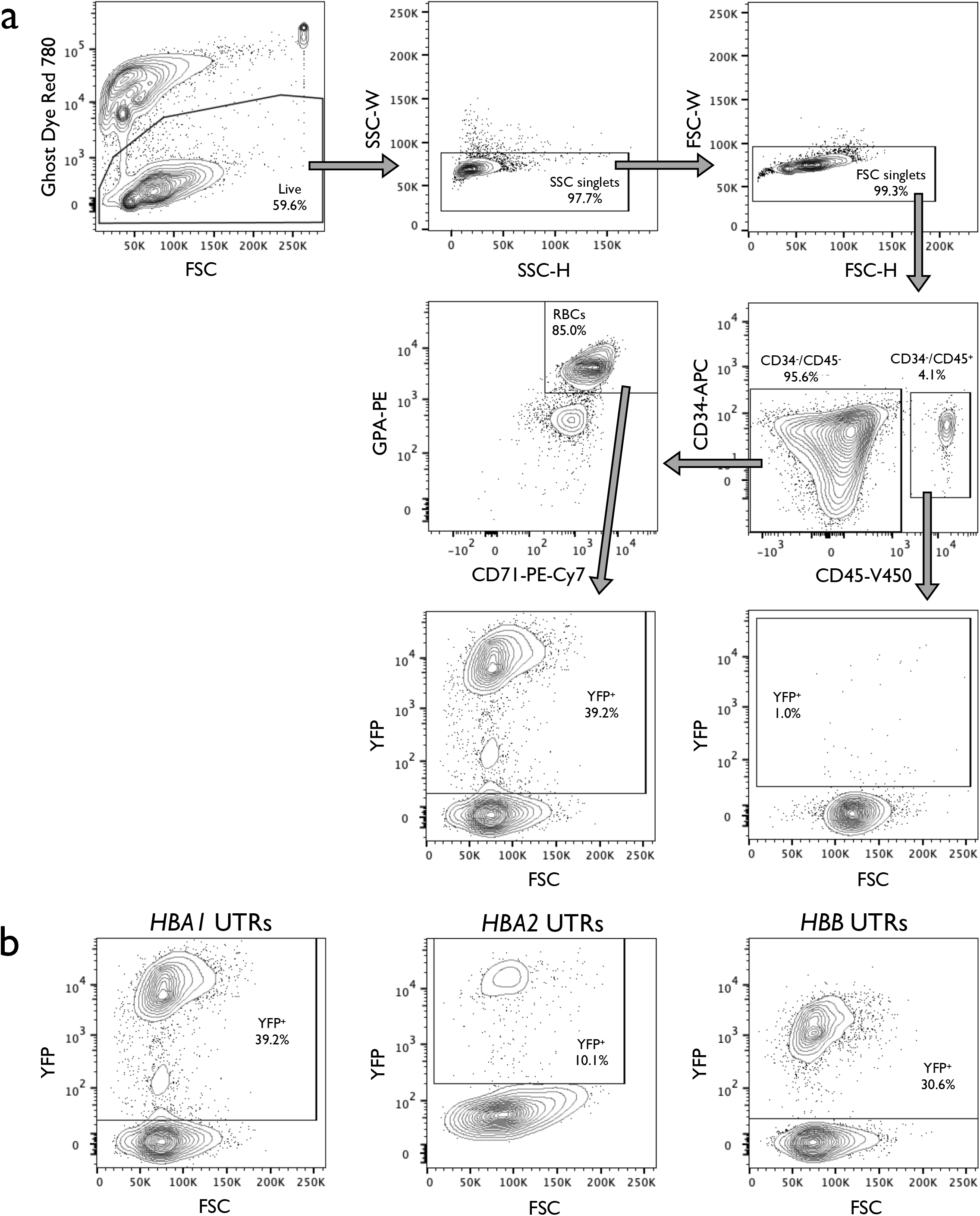
Representative staining and gating scheme used to analyze targeting and differentiation rates of RBCs. A) Representative flow cytometry staining and gating scheme for human HSPCs targeted at *HBA1* with HBB-T2A-YFP (*HBA1* UTRs) and differentiated into RBCs over the course of a 14-day protocol. This indicates that only RBCs (CD34^−^/CD45^−^/CD71^+^/GPA^+^) are able to express the integrated T2A-YFP marker. Analysis was performed on BD FACS Aria II platform. B) Representative YFP x FSC flow cytometry images of of RBCs (CD34^−^/CD45^−^/CD71^+^/GPA^+^) derived from HSPCs targeted with *HBA1* UTRs, *HBA2* UTRs, and *HBB* UTRs vector. AAV only controls were used for each vector to establish gating scheme, leading to slight variation in positive/negative cut-offs across images.

**Supplemental Figure 7:**
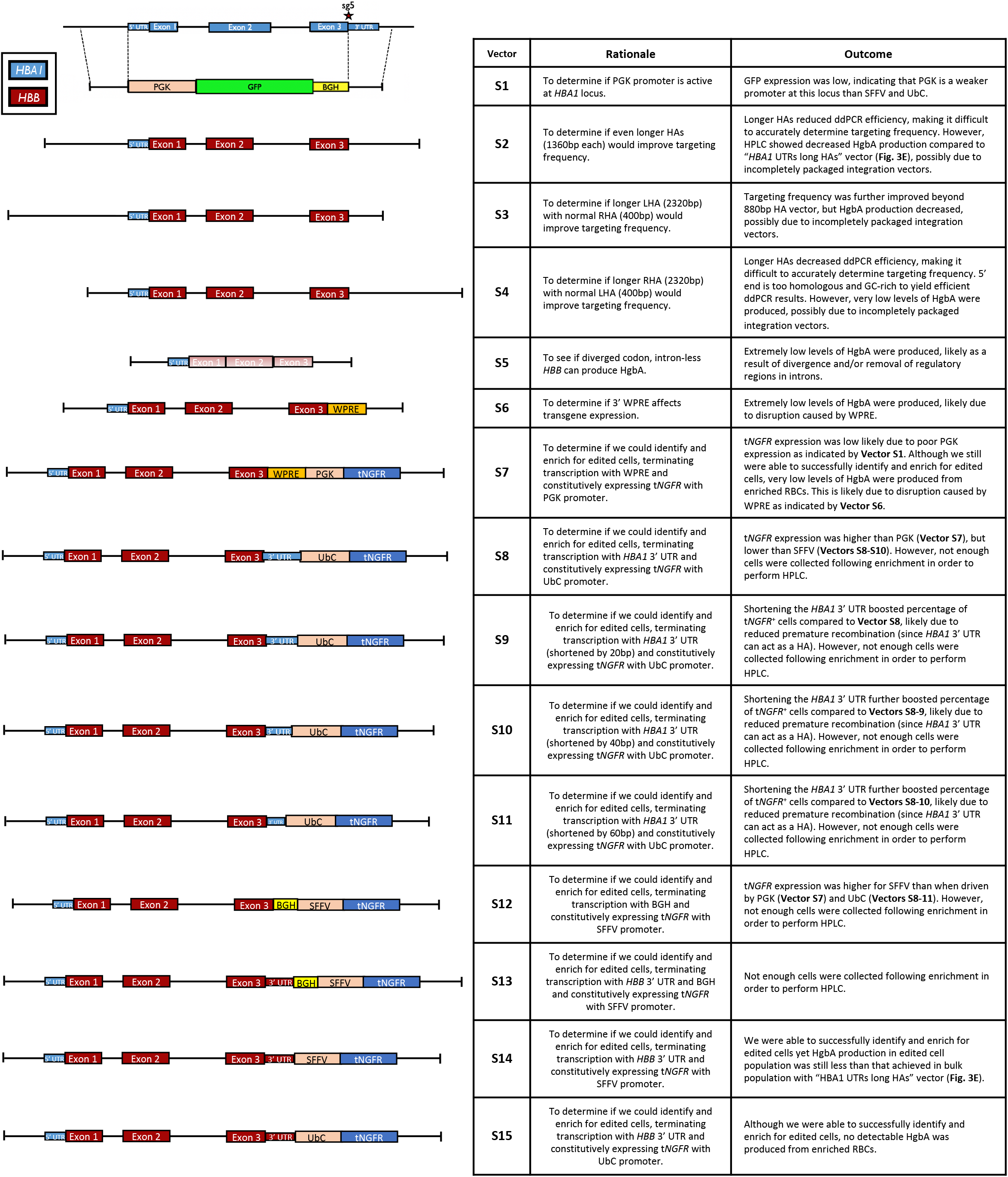
Integration cassettes screened for development of clinical vector. Displayed are schematics and corresponding rationale for design as well as eventual outcomes for Vectors S1-6.

**Supplemental Figure 8:**
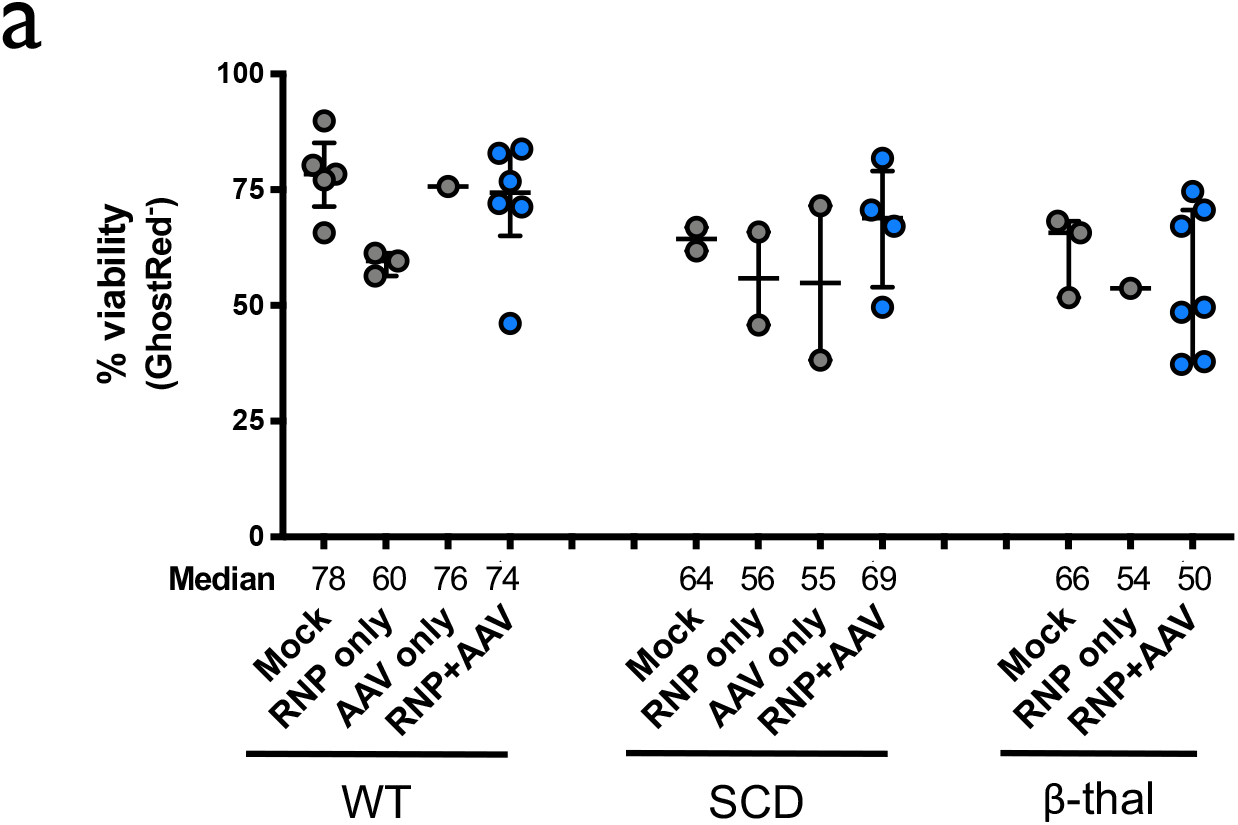
Viability of HSPCs post-editing. HSPC viability was quantified 2-4d post-editing by flow cytometry. Depicted are the percentage of cells that stained negative for GhostRed viability dye. All cells were edited with our optimized HBB gene replacement vector using standard conditions (i.e. electroporation of Cas9 RNP+sg5, 5K MOI of AAV, and no AAV wash at 24h). Bars represent median + interquartile range. WT: N = 5 for mock, N = 3 for RNP only, N = 1 for AAV only, and N = 6 for RNP+AAV treatment group; SCD: N = 2 for each treatment group with exception of RNP+AAV with N =4; β-thal: N = 3 for mock, N = 1 for RNP only, and N = 7 for RNP+AAV treatment group.

**Supplemental Figure 9:**
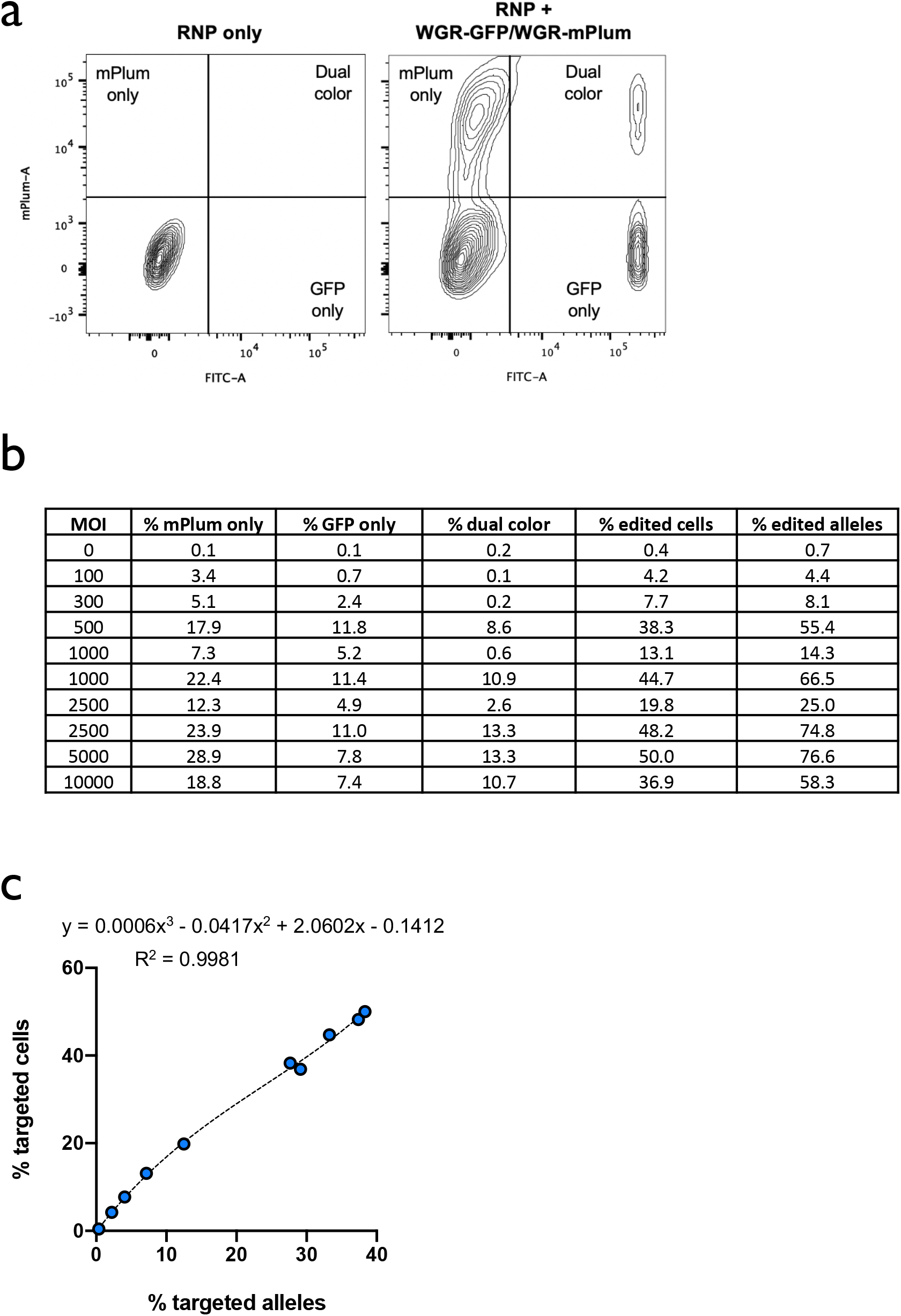
Relationship between % edited alleles and % edited cells. A) Representative FACS plots of CD34+ HSPCs simultaneously targeted by HBA1-WGR-GFP AAV6 (shown in Fig. 1c) and HBA1-WGR-mPlum AAV6. B) Table showing % of populations targeted with GFP only, mPlum only, and both colors. Percent of edited cells was then converted to % edited alleles by the following equation: (total % targeted cells + (% dual color)*2)/2 = total % targeted alleles. C) Percent edited cells is plotted against % edited alleles for data shown in panel B. A polynomial regression (R^2^ = 0.9981)was used to determine an equation to convert between the % edited alleles to % edited cells and vice versa.

**Supplemental Figure 10:**
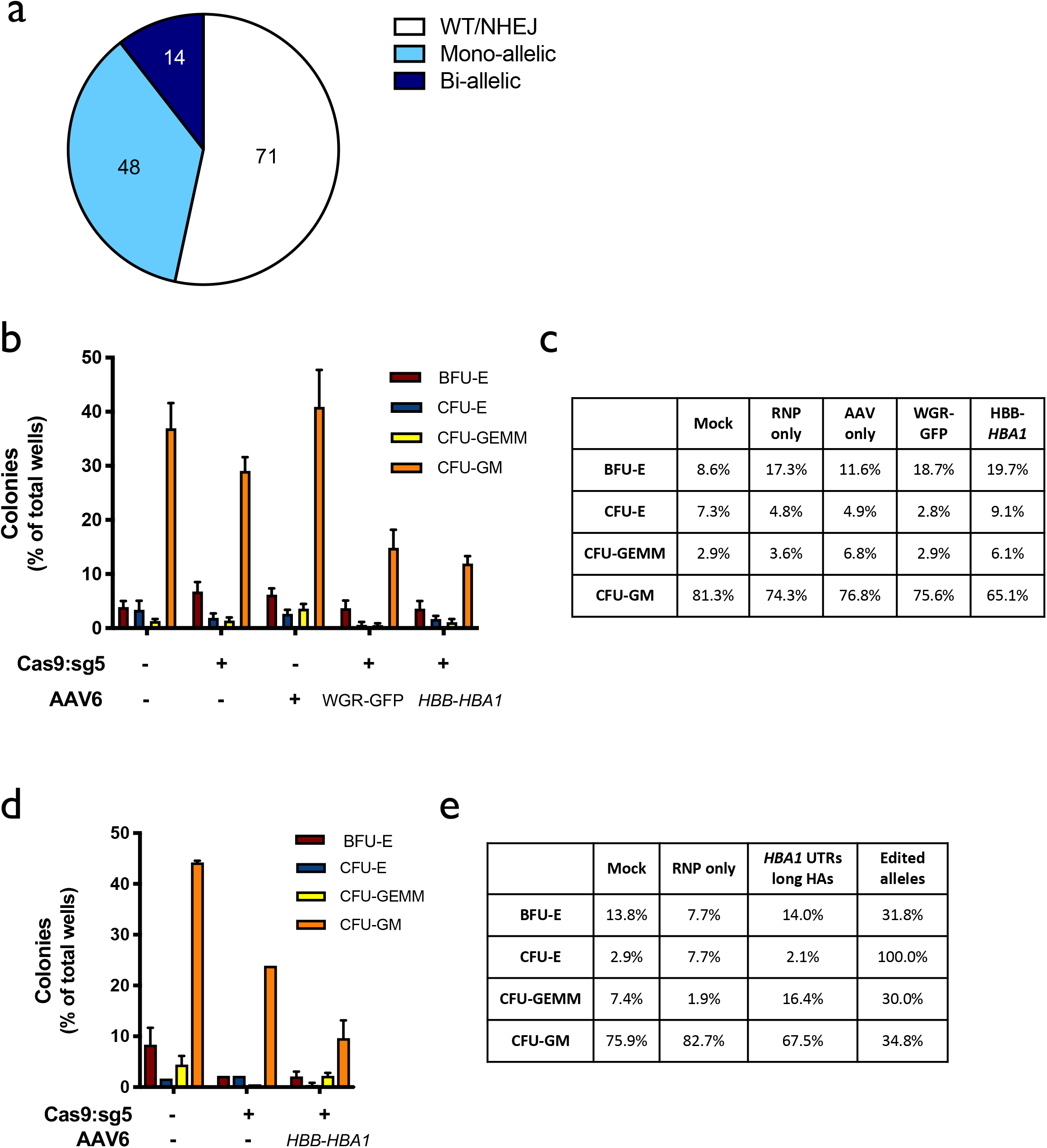
Analysis of colony-forming units of HSPCs plated into methylcellulose. A) Distribution of genotypes of methylcellulose colonies displayed in Panels B and D. Numbers of clones corresponding to each category are included in the pie chart. B) *In vitro* (pre-engraftment) live CD34^+^ HSPCs from healthy donors were single-cell sorted into 96-well plates containing semisolid methylcellulose media for colony forming assays. 14d post-sorting cells were analyzed for morphology. Depicted are number of colonies formed for each lineage (CFU-E = erythroid lineage; CFU-GEMM = multi-lineage; or CFU-GM = granulocyte/macrophage lineage) divided by the total number of wells available for colonies. N=2 experimental replicates with a minimum of 3 96-well methylcellulose-coated plates for each treatment. N = 3 experimental replicates with a minimum of 3 96-well methylcellulose-coated plates for mock, RNP only, and WGR-GFP AAV6 treatment groups; N = 2 for AAV only and *HBB-HBA1* AAV6 treatment groups. C) Percent distribution of each lineage among all colonies for each treatment for Panel B. D) *In vitro* (pre-engraftment) live CD34^+^ β-thalassemia patient-derived HSPCs were singlecell sorted into 96-well plates containing semisolid methylcellulose media for colony forming assays. 14d post-sorting cells were analyzed for morphology. Depicted are number of colonies formed for each lineage (B = BFU-E and C = CFU-E (erythroid lineage); GE = CFU-GEMM (multi-lineage); or GM = CFU-GM (granulocyte/macrophage lineage)) divided by the total number of wells available for colonies. For Mock and RNP+AAV, N=2 experimental replicates with a minimum of 3 96-well methylcellulose-coated plates for each treatment; N=1 experimental replicate with 3 plates for RNP only treatment. E) Percent distribution of each lineage among all colonies for each treatment for Panel.

**Supplemental Figure 11:**
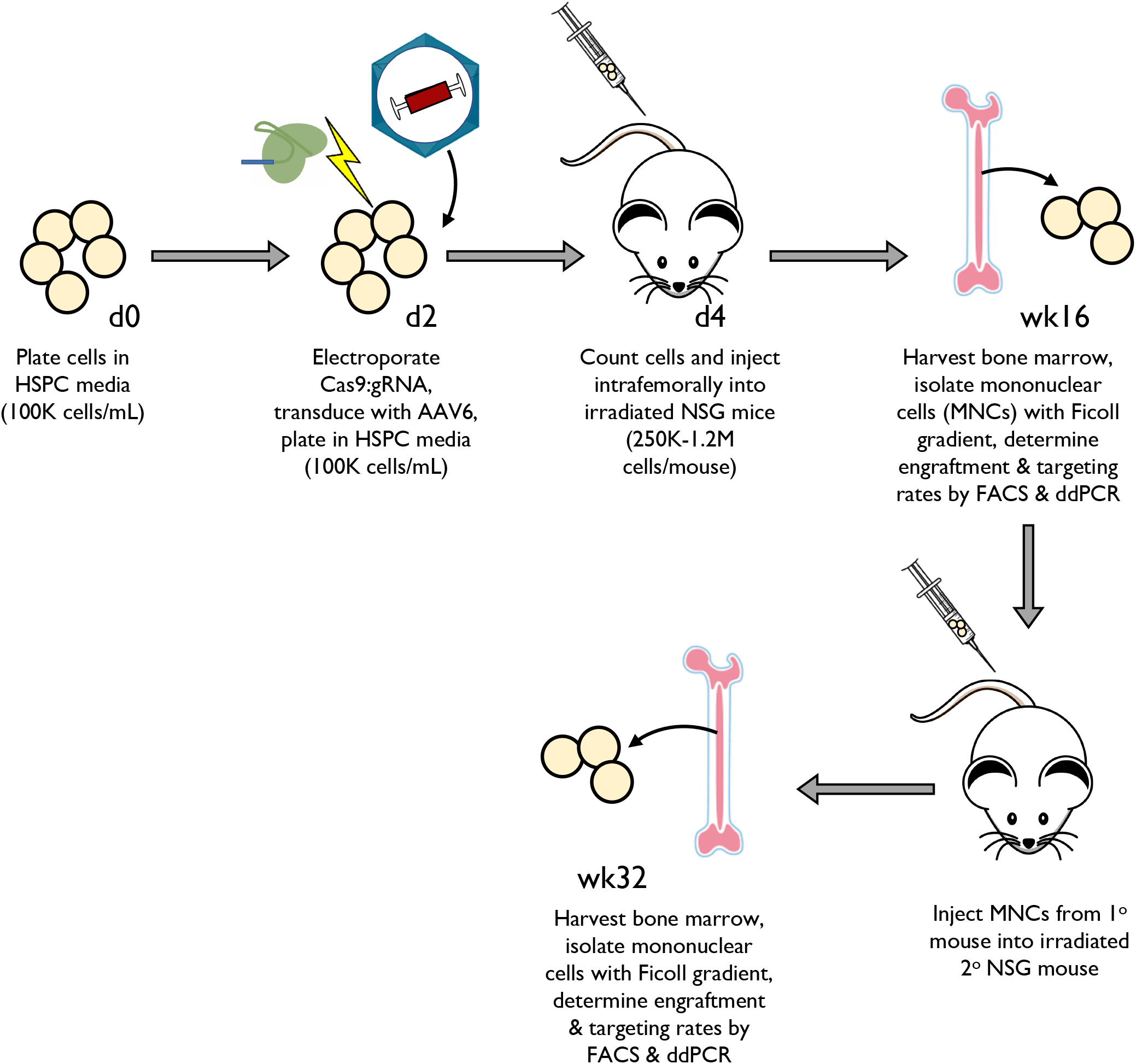
Timeline for targeting of HSPCs and transplantation into mice. Timeline for targeting of HSPCs with HBB integration vectors, transplantation into mice (both 1° and 2° engraftment), and subsequent analysis.

**Supplemental Figure 12:**
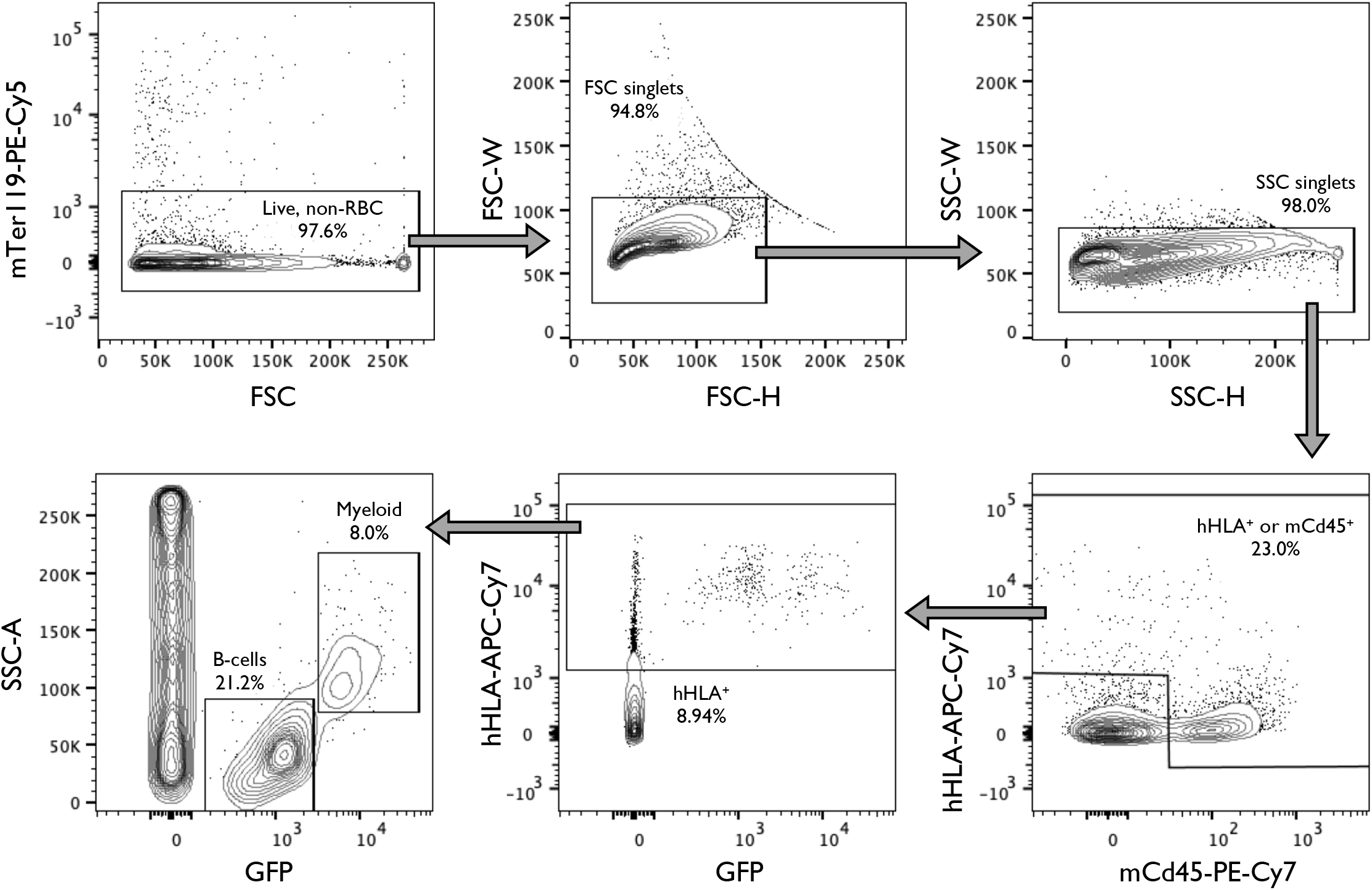
Representative staining and gating scheme used to analyze engraftment and targeting rates of human HSPCs into NSG mice. Representative flow cytometry staining and gating scheme used to analyze targeting and engraftment rates of human HSPCs transplanted into the bone marrow of NSG mice. This sample was targeted with a UbC-GFP integration at the *HBA1* locus. This demonstrates that only human cells (hHLA^+^) are able to express GFP. Analysis was performed on BD FACS Aria II platform.

**Supplemental Figure 13:**
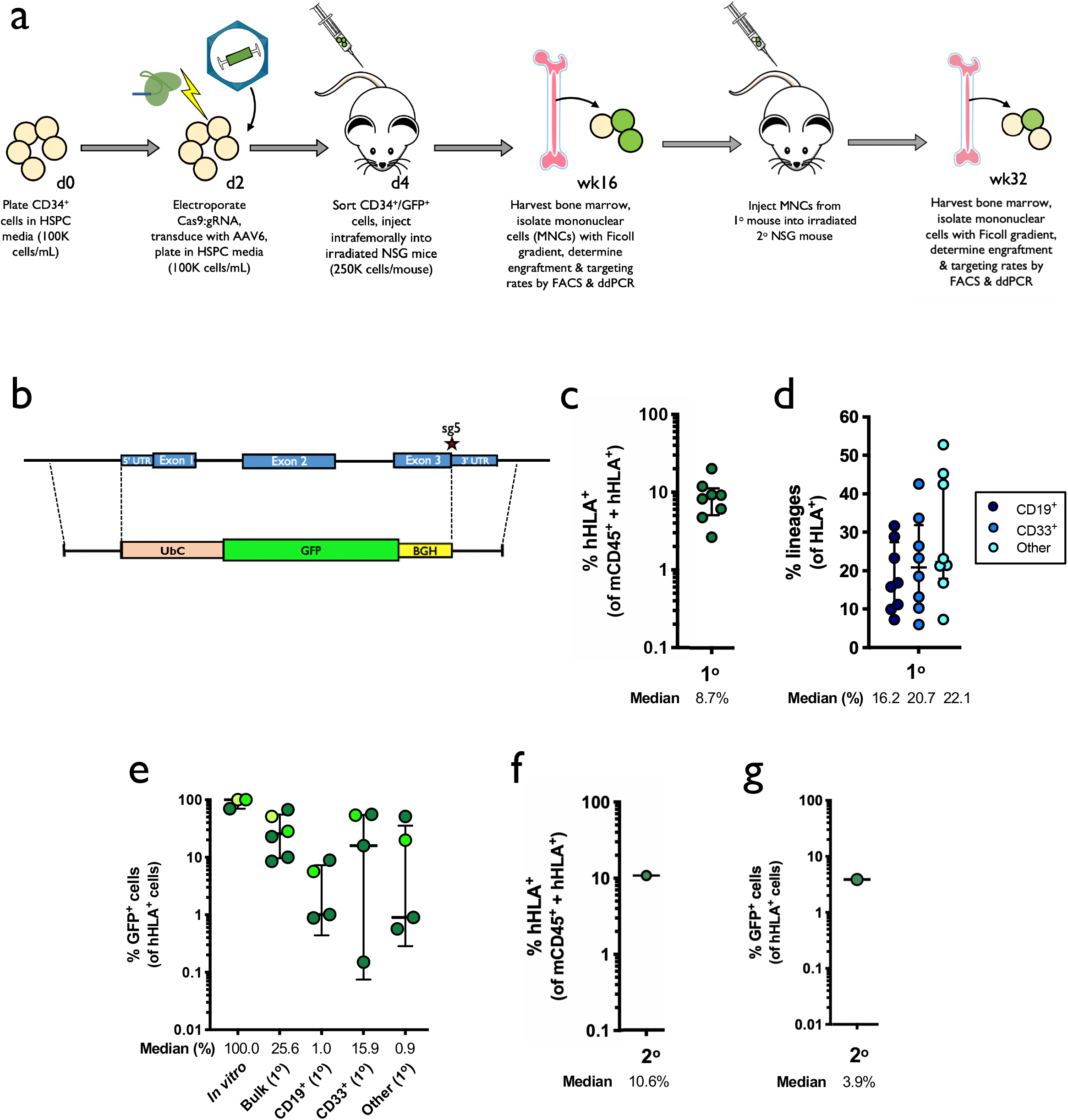
Engraftment of human HSPCs targeted with GFP at α-globin locus into NSG mice. A) Timeline for targeting of HSPCs with UbC-GFP integration vector, transplantation into mice (both 1° and 2° engraftment), and subsequent analysis. B) AAV6 DNA repair donor design schematic to introduce a UbC-GFP-BGH integration is depicted at the *HBA1* locus. C) 16 weeks after bone marrow transplantation of targeted human CD34^+^ HSPCs into NSG mice, bone marrow was harvested and rates of engraftment were determined (1°). Depicted is the percentage of mTerr119^−^ cells (non-RBCs) that were hHLA^+^ from the total number of cells that were either mCd45^+^ or hHLA^+^. Bars represent median + interquartile range. N = 8. D) Among engrafted human cells, the distribution among CD19^+^ (B-cell), CD33^+^ (myeloid), or other (i.e. HSPC/RBC/T/NK/Pre-B) lineages are indicated. Bars represent median + interquartile range. N = 8. E) Percentage of GFP^+^ cells among pre-transplantation (*in vitro*, post-sorting) and successfully-engrafted populations, both bulk HSPCs and among CD19^+^ (B-cell), CD33^+^ (myeloid), and other lineages. Bars represent median + interquartile range. N > 3 for each treatment group. F) Following primary engraftments, engrafted human cells were transplanted a second time into the bone marrow of NSG mice. 16 weeks post-transplantation, bone marrow was harvested and rates of engraftment were determined (2°). Depicted is the percentage of mTerr119^−^ cells (non-RBCs) that were hHLA^+^ from the total number of cells that were either mCd45^+^ or hHLA^+^. N = 1. G) Percentage of GFP^+^ cells among successfully-engrafted population from the secondary transplant depicted in Panel F. N = 1.

**Supplemental Figure 14:**
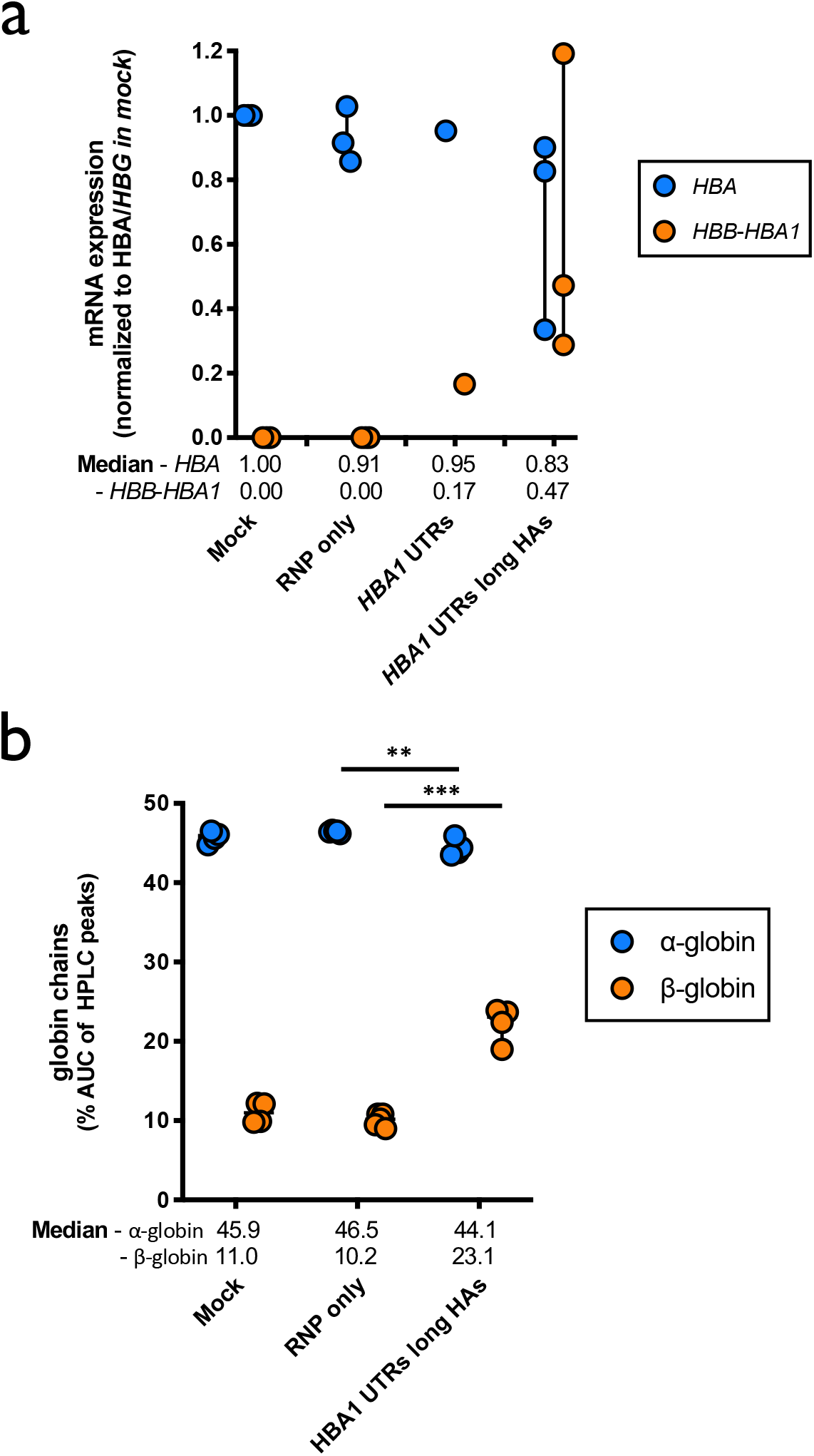
Characterization of targeted β-thalassemia-derived HSPCs. A) Following differentiation of targeted HSPCs into RBCs, mRNA was harvested and converted into cDNA. Expression of HBA (does not distinguish between *HBA1* and *HBA2*) and *HBB* transgene were normalized to *HBG* expression. Bars represent median + interquartile range. N = 3 for each treatment group with exception of *HBA1* UTRs with N = 1. B) Summary of reverse-phase globin chain HPLC results showing % AUC of β-globin and α-globin. Bars represent median + interquartile range. Bars represent median + interquartile range. **: P<0.005; ***: P<0.0001 determined using unpaired t test. N > 4 for each treatment group.

## METHODS

### AAV6 vector design, production, and purification

All AAV6 vectors were cloned into the pAAV-MCS plasmid (Agilent Technologies, Santa Clara, CA, USA), which contains inverted terminal repeats (ITRs) derived from AAV2. Gibson Assembly Mastermix (New England Biolabs, Ipswich, MA, USA) was used for the creation of each vector as per manufacturer’s instructions. Cut site (CS) vectors were designed such that the left and right homology arms (“LHA” and “RHA”, respectively) are immediately flanking the cut site at either *HBA2* or *HBA1* gene. Whole gene replacement (WGR) vectors have a LHA flanking the 5’ UTR of either the *HBA2* or *HBA1* gene while the RHA immediately flanks downstream of its corresponding cut site. The LHA and RHA of every vector is 400bp, unless otherwise noted, with the vector name (*HBA2/HBA1* and CS/WGR) referencing the target integration type and homology arms used, respectively. Within Figure 1, CS and WGR vectors consisted of a SFFV-GFP-BGH expression cassette. An alternative promoter, UbC, was also used in creating a WGR vector for *HBA1* **(Supplemental Fig. 10)**. In Figure 2, WGR-T2A-YFP vectors consisted of the full-length *HBB* gene, unless noted, with a T2A-YFP expression cassette immediately following exon 3 of the *HBB* gene using the LHA and RHA described previously for WGR. These full-length HBB-T2A-YFP vectors were either flanked by 5’ and 3’ UTRs of HBB, *HBA2*, or *HBA1* as denoted in Figure 2a. In subsequent experiments, for targeting of SCD or β-thalassemia patient-derived CD34^+^ HSPCs, WGR vectors were designed to target the *HBA1* site and contained a full-length *HBB* gene flanked by either *HBA1* UTRs or *HBB* UTRs. While the *‘HBB* UTRs’ and *‘HBA1* UTRs’ vector both share 400bp HAs, the *‘HBA1* UTRs long HAs’ vector was modified to have 880bp HAs. Few modifications were made to the production of AAV6 vectors as described^41^. 293T cells (Life Technologies, Carlsbad, CA, USA) were seeded in ten 15 cm^2^ dishes with 13-15×10^6^ cells per plate. 24h later, each dish was transfected with a standard polyethylenimine (PEI) transfection of 6μg ITR-containing plasmid and 22μg pDGM6 (gift from David Russell, University of Washington, Seattle, WA, USA), which contains the AAV6 cap genes, AAV2 rep genes, and Ad5 helper genes. After a 48-72h incubation, cells were lysed by 3 freeze-thaw cycles, treated with benzonase (Thermo Fisher Scientific, Waltham, MA, USA) at 250 U/mL, and the vector was then purified through an iodixanol gradient centrifugation at 48,000 RPM for 2.25h at 18°C. Afterwards, full capsids were isolated at the 40–58% iodixanol interface and then stored at 80°C until further use. As an alternative method, AAVPro Purification Kit (All Serotypes)(Takara Bio USA, Mountain View, CA, USA) were also used following the 48-72 h incubation period, to extract full AAV6 capsids as per manufacturer’s instructions. AAV6 vectors were titered using ddPCR to measure number of vector genomes as previously described^42^.

### Culturing of CD34^+^ HSPCs

Human CD34^+^ HSPCs were cultured as previously described^18,24,36,39,43,44^. CD34^+^ HSPCs were sourced from fresh cord blood (generously provided by Binns Family program for Cord Blood Research), frozen cord blood and Plerixafor- and/or G-CSF-mobilized peripheral blood (AllCells, Alameda, CA, USA and STEMCELL Technologies, Vancouver, Canada), frozen Plerixafor- and/or G-CSF-mobilized peripheral blood of patients with SCD, and frozen Plerixafor- and Filgrastim-mobilized peripheral blood from β-thalassemia patient (compound heterozygous - c.-138C>T and c.92+5G>C). The β-thalassemia-derived HSPCs were collected under protocol 14-H-0077, (registered on clinicaltrials.gov, NCT02105766), which was approved and renewed annually by the NHLBI IRB. The patient provided informed consent for the study. CD34^+^ HSPCs were cultured at 2.5×10^5^–5×10^5^ cells/mL in StemSpan SFEM II (STEMCELL Technologies, Vancouver, Canada) base medium supplemented with stem cell factor (SCF)(100ng/mL), thrombopoietin (TPO)(100ng/mL), FLT3–ligand (100ng/mL), IL-6 (100ng/mL), UM171 (35nM), 20mg/mL streptomycin, and 20U/mL penicillin. The cell incubator conditions were 37°C, 5% CO2, and 5% O2.

### Genome editing of CD34^+^ HSPCs

Chemically-modified sgRNAs used to edit CD34^+^ HSPCs at either *HBA2* or *HBA1* were purchased from Synthego (Menlo Park, CA, USA) and TriLink BioTechnologies (San Diego, CA, USA) and were purified by high-performance liquid chromatography (HPLC). The sgRNA modifications added were the 2’-O-methyl-3’-phosphorothioate at the three terminal nucleotides of the 5’ and 3’ ends described previously^33^. The target sequences for sgRNAs were as follows: *sg1*: 5’-CTACCGAGGCTCCAGCTTAA-3’; *sg2*: 5’-GGCAGGAGGAACGGCTACCG-3’; *sg3*: 5’-GGGGAGGAGGGCCCGTTGGG-3’; *sg4*: 5’-CCACCGAGGCTCCAGCTTAA-3’; and *sg5*: 5’-GGCAAGAAGCATGGCCACCG-3’. All Cas9 protein (Alt-R S.p. Cas9 Nuclease V3) used was purchased from Integrated DNA Technologies (Coralville, Iowa, USA). The RNPs were complexed at a Cas9: sgRNA molar ratio of 1:2.5 at 25°C for 10min prior to electroporation. CD34^+^ cells were resuspended in P3 buffer (Lonza, Basel, Switzerland) with complexed RNPs and electroporated using the Lonza 4D Nucleofector (program DZ-100). Cells were plated at 2.5×10^5^ cells/mL following electroporation in the cytokine-supplemented media described previously. Immediately following electroporation, AAV6 was supplied to the cells at 5×10^3^-1×10^4^ vector genomes/cell based on titers determined by ddPCR.

### Indel frequency analysis by TIDE

2-4d post-targeting, HSPCs were harvested and QuickExtract DNA extraction solution (Epicentre, Madison, WI, USA) was used to collect gDNA. The following primers were then used to amplify respective cut sites at *HBA2* and *HBA1* along with CleanAmp PCR 2x Master Mix (TriLink, San Diego, CA, USA) according to manufacturer’s instructions: *HBA2* (sg1-3): forward: 5’-CCCGAAAGGAAAGGGTGGCG-3’ reverse: 5’-TGGCACCTGCACTTGCACTG-3’; *HBA1* (sg4-5): forward: 5’-TCCGGGGTGCACGAGCCGAC-3’, reverse: 5’-GCGGTGGCTCCACTTTCCCT-3’. PCR reactions were then run on a 1% agarose gel and appropriate bands were cut and gel-extracted using a GeneJET Gel Extraction Kit (Thermo Fisher Scientific, Waltham, MA, USA) according to manufacturer’s instructions. Gel-extracted amplicons were then Sanger sequenced with the following primers: *HBA2* (sg1-3): forward: 5’-GGGGTGCGGGCTGACTTTCT-3’ reverse: 5’-CTGAGACAGGTAAACACCTCCAT-3’; *HBA1* (sg4-5): forward: 5’-TGGAGACGTCCTGGCCCC-3’, reverse: 5’-CCTGGCACGTTTGCTGAGG-3’. Resulting Sanger chromatograms were the used as input for indel frequency analysis by TIDE as previously described^34^.

### Gene targeting analysis by flow cytometry

4-8d post-targeting with fluorescent gene replacement vectors, CD34^+^ HSPCs were harvested and the percentage of edited cells was determined by flow cytometry. Cells were analyzed for viability using Ghost Dye Red 780 (Tonbo Biosciences, San Diego, CA, USA) and reporter expression was assessed using either the Accuri C6 flow cytometer (BD Biosciences, San Jose, CA, USA) or FACS Aria II (BD Biosciences, San Jose, CA, USA). The data was subsequently analyzed using FlowJo (FlowJo LLC, Ashland, OR, USA).

### Allelic targeting analysis by ddPCR

2-4d post-targeting, HSPCs were harvested and QuickExtract DNA extraction solution (Epicentre, Madison, WI, USA) was used to collect gDNA. gDNA was then digested using BAMH1-HF as per manufacturer’s instructions (New England Biolabs, Ipswich, MA, USA). The percentage of targeted alleles within a cell population was measured by ddPCR using the following reaction mixture: 1-4μL of digested gDNA input, 10μL ddPCR SuperMix for Probes (No dUTP)(Bio-Rad, Hercules, CA, USA), primer/probes (1:3.6 ratio; Integrated DNA Technologies, Coralville, Iowa, USA), volume up to 20μL with H_2_O. ddPCR droplet were then generated following the manufacturer’s instructions (Bio-Rad, Hercules, CA, USA): 20μL of ddPCR reaction, 70μL droplet generation oil, and 40μL of droplet sample. Thermocycler (Bio-Rad, Hercules, CA, USA) settings were as follows: 1. 98°C (10min), 2. 94°C (30s), 3. 57.3°C (30s), 4. 72°C (1.75min)(return to step 2 × 40–50 cycles), 5. 98°C (10 min). Analysis of droplet samples was done using the QX200 Droplet Digital PCR System (Bio-Rad, Hercules, CA, USA). To determine percentage of alleles targeted, the number of Poisson-corrected integrant copies/mL were divided by the number of Poisson-corrected reference DNA copies/mL. The following primers and 6-FAM/ZEN/IBFQ-labelled hydrolysis probes were purchased as custom-designed PrimeTime qPCR Assays from Integrated DNA Technologies (Coralvilla, IA, USA): All *HBA2*-GFP vectors (spans from BGH to outside 400bp *HBA2* RHA): forward: 5’-TAGTTGCCAGCCATCTGTTG-3’, reverse: 5’-GGGGACAGCCTATTTTGCTA-3’, probe: 5’-AAATGAGGAAATTGCATCGC-3’; All *HBA1*-GFP vectors (spans from BGH to outside 400bp *HBA1* RHA): forward: 5’-TAGTTGCCAGCCATCTGTTG-3’, reverse: 5’-TAGTGGGAACGATGGGGGAT-3’, probe: 5’-AAATGAGGAAATTGCATCGC-3’; *HBA2*-HBB-T2A-YFP vector (spans from YFP to outside 400bp *HBA2* RHA): forward: 5’-AGTCCAAGCTGAGCAAAGA-3’, reverse: 5’-GGGGACAGCCTATTTTGCTA-3’, probe: 5’-CGAGAAGCGCGATCACATGGTCCTGC-3’; All *HBA1*-HBB-T2A-YFP vectors (spans from YFP to outside 400bp *HBA1* RHA): forward: 5’-AGTCCAAGCTGAGCAAAGA-3’, reverse: 5’-TAGTGGGAACGATGGGGGAT-3’, probe: 5’-CGAGAAGCGCGATCACATGGTCCTGC-3’; *HBA1*-*HBB* vectors (with 400bp HAs, without T2A-YFP)(spans from *HBB* exon 3 to outside 400bp *HBA1* RHA): forward: 5’-GCTGCCTATCAGAAAGTGGT-3’, reverse: 5’-TAGTGGGAACGATGGGGGAT-3’, probe: 5’-CTGGTGTGGCTAATGCCCTGGCCC-3’; *HBA1-HBB* vector (with 880bp HAs, without T2A-YFP)(spans from *HBB* exon 3 to outside 880bp *HBA1* RHA): forward: 5’-GCTGCCTATCAGAAAGTGGT-3’, reverse: 5’-ATCACAAACGCAGGCAGAG-3’, probe: 5’-CTGGTGTGGCTAATGCCCTGGCCC-3’. The primers and HEX/ZEN/IBFQ-labelled hydrolysis probe purchased as custom-designed PrimeTime qPCR Assays from Integrated DNA Technologies (Coralvilla, IA, USA) were used to amplify the *CCRL2* reference gene: forward: 5’-GCTGTATGAATCCAGGTCC-3’, reverse: 5’-CCTCCTGGCTGAGAAAAAG-3’, probe: 5’-TGTTTCCTCCAGGATAAGGCAGCTGT-3’. Due to the length of the ‘HBA1 UTRs long HAs’ vector and to ensure episomal AAV is not detected, the ddPCR amplicon exceeds the template size recommended by the ddPCR manufacturer. Upon analysis of the data, the percentage of targeted alleles of this vector is underestimated. Therefore, in these instances a correction factor to account for this underestimation was determined by amplifying gDNA harvested from HSPCs targeted with *HBA1* UTRs vector with 400bp HAs with both sets of ddPCR primers and probes (those for vectors with 400bp and 880bp HAs) as well as *CCRL2* reference probes. The resulting correction factor was then applied to the targeted allele percentage from samples targeted with and amplified with primers and probe for 880bp HAs.

### Off-target activity analysis by rhAmpSeq

Predicted off-target sites for *HBA1* sg5 was identified using COSMID with up to three mismatches allowed in the 19 PAM-proximal bases and the PAM sequence NGG. rhAmpSeq targeted sequencing was performed for the 40 most highly-predicted off-target sites as described previously.

### In vitro differentiation of CD34^+^ HSPCs into erythrocytes

Following targeting, HSPCs derived from healthy, SCD, or β-thalassemia patients were cultured for 14-16d at 37°C and 5% CO2 in SFEM II medium (STEMCELL Technologies, Vancouver, Canada) as previously described^37,38^. SFEMII base medium was supplemented with 100U/mL penicillin–streptomycin, 10ng/mL SCF, 1ng/mL IL-3 (PeproTech, Rocky Hill, NJ, USA), 3U/mL erythropoietin (eBiosciences, San Diego, CA, USA), 200μg/mL transferrin (Sigma-Aldrich, St. Louis, MO, USA), 3% antibody serum (heat-inactivated from Atlanta Biologicals, Flowery Branch, GA, USA), 2% human plasma (umbilical cord blood), 10μg/mL insulin (Sigma-Aldrich, St. Louis, MO, USA) and 3U/mL heparin (Sigma-Aldrich, St. Louis, MO, USA). In the first phase, d 0-7 (day zero being 2d posttargeting) of differentiation, cells were cultured at 1×10^5^ cells/mL. In the second phase, d7–10, cells were maintained at 1×10^5^ cells/mL, and IL-3 was removed from the culture. In the third phase, d11–16, cells were cultured at 1×10^6^ cells/mL, and transferrin was increased to 1 mg/mL within the culture medium.

### mRNA analysis

Following differentiation of HSPCs into erythrocytes, cells were harvested and RNA was extracted using RNeasy Plus Mini Kit (Qiagen, Hilden, Germany). Subsequently, cDNA was made from approximately 100ng RNA using the iScript Reverse Transcription Supermix for RT-qPCR (Bio-Rad, Hercules, CA, USA). Expression levels of β-globin transgene and α-globin mRNA were quantified by ddPCR using the following primers and 6-FAM/ZEN/IBFQ-labelled hydrolysis probes purchased as custom-designed PrimeTime qPCR Assays from Integrated DNA Technologies (Coralvilla, IA, USA): *HBB*: forward: 5’-GAGAACTTCAGGCTCCTG-3’, reverse: 5’-CGGGGGTACGGGTGCAGGAA-3’, probe: 5’-TGGCCATGCTTCTTGCCCCT-3’; *HBA* (does not distinguish between *HBA2* and *HBA1)*: forward: 5’-GACCTGCACGCGCACAAGCTT-3’, reverse: 5’-GCTCACAGAAGCCAGGAACTTG-3’, probe: 5’-CAACTTCAAGCTCCTAAGCCA-3’. To normalize for RNA input, levels of the RBC-specific reference gene *GPA* was determined in each sample using the following primers and HEX/ZEN/IBFQ-labelled hydrolysis probes purchased as custom-designed PrimeTime qPCR Assays from Integrated DNA Technologies (Coralvilla, IA, USA): forward: 5’-ATATGCAGCCACTCCTAGAGCTC-3’, reverse: 5’-CTGGTTCAGAGAAATGATGGGCA-3’, probe: 5’-AGGAAACCGGAGAAAGGGTA-3’. ddPCR reactions were created using the respective primers and probes and droplets were generated as described above. Thermocycler (Bio-Rad, Hercules, CA, USA) settings were as follows: 1. 98°C (10min), 2. 94°C (30s), 3. 59.4°C (30s), 4. 72°C (30s)(return to step 2 × 40–50 cycles), 5. 98°C (10 min). Analysis of droplet samples was done using the QX200 Droplet Digital PCR System (Bio-Rad, Hercules, CA, USA). To determine relative expression levels, the number of Poisson-corrected HBA or HBB transgene copies/mL were divided by the number of Poisson-corrected GPA copies/mL.

### Immunophenotyping of differentiated erythrocytes

HSPCs subjected to the above erythrocyte differentiation were analyzed at d14-16 for erythrocyte lineage-specific markers using a FACS Aria II (BD Biosciences, San Jose, CA, USA). Edited and non-edited cells were analyzed by flow cytometry using the following antibodies: hCD45 V450 (HI30; BD Biosciences, San Jose, CA, USA), CD34 APC (561; BioLegend, San Diego, CA, USA), CD71 PE-Cy7 (OKT9; Affymetrix, Santa Clara, CA, USA), and CD235a PE (GPA)(GA-R2; BD Biosciences, San Jose, CA, USA).

### Steady-state hemoglobin tetramer analysis

HSPCs subjected to the above erythrocyte differentiation were lysed using water equivalent to three volumes of pelleted cells. The mixture was incubated at room temperature for 15min, followed by 30s sonication. For separation of lysate from the erythrocyte ghosts, centrifugation was performed at 13,000 RPM for 5min. HPLC analysis of hemoglobins in their native form were analyzed on a weak cation-exchange PolyCAT A column (100 × 4.6-mm, 3μm, 1,000Å) (PolyLC Inc., Columbia, MD, USA) using a Shimadzu UFLC system at room temperature. Mobile phase A (MPA) consists of 20mM Bis-tris + 2mM KCN, pH 6.96. Mobile phase B (MPB) consists of 20mM Bis-tris + 2mM KCN + 200mM NaCl, pH 6.55. Clear hemolysate was diluted four times in MPA, and then 20μL was injected onto the column. A flow rate of 1.5mL/min and the following gradients were used in time (min)/%B organic solvent: (0/10%; 8/40%; 17/90%; 20/10%; 30/stop).

### Reverse-phase HPLC globin chain analysis

Analysis of globin chains in CD34^+^ cell-derived erythroblasts was performed by reverse-phase HPLC, as previously described^45,46^. In brief, the reverse-phase HPLC assay was carried out on an Agilent 1260 Infinity II HPLC system with Diode-Array Detector. The chromatographic column is Aeris™ 3.6μm WIDEPORE XB-C18 200Å, LC Column 250 × 4.6mm behind a securityGuard™ ULTRA cartridge (Phenomenex). Globin chains were separated using a gradient program of 41–47% solvent B (acetonitrile) mixing with solvent A (0.1% trifluoroacetic acid in HPLC grade water at pH 2.9) and quantified by the area under the curve of the corresponding peaks in reverse-phase HPLC chromatogram.

### Methylcellulose CFU assessment

2d post-targeting, HSPCs were stained using CD34 APC (561; BioLegend, San Diego, CA, USA), Ghost Dye Red 780 (Tonbo Biosciences, San Diego, CA, USA) and live CD34^+^ cells were sorted into 96-well plates containing MethoCult Optimum (STEMCELL Technologies, Vancouver, Canada). After 12–16d, colonies were appropriately scored based on external appearance in a blinded fashion.

### CD34^+^ HSPC transplantation into immunodeficient NSG mice

Six-to eight-week-old NSG mice (Jackson Laboratory, Bar Harbor, ME, USA) were irradiated using 200rads of radiation 12-24h prior to transplantation with targeted HSPCs (2d post-targeting) via intrafemoral or tail-vein injections. Approximately 2.5×10^5^-1.3×10^6^ electroporated HSPCs (exact number noted in figures) were injected using an insulin syringe with a 27G, 0.5 inch (12.7mm) needle. This experimental protocol was approved by Stanford University’s Administrative Panel on Laboratory Animal Care. All mouse studies reported in this paper were performed as a minimum of three separate experimental replicates of editing and transplantation. For sample size, we transplanted as many mice as was feasible to cover the non-Gaussian distribution that would be expected from experimental and donor variability, while also minimizing the total number of animals as per FDA’s Center for Biologics Evaluation and Research guidelines.

### Assessment of Human Engraftment

15-17wks post-transplantation of CD34^+^-edited HSPCs, mice were euthanized and bone marrow was harvested from tibia, femurs, pelvis, sternum, and spine using a pestle and mortar. Mononuclear cells were enriched using a Ficoll gradient centrifugation (Ficoll-Paque Plus, GE Healthcare, Chicago, IL) for 25min at 2,000g at room temperature. The samples were then stained for 30min at 4°C with the following antibodies: monoclonal anti-human CD33 V450 (WM53; BD Biosciences, San Jose, CA, USA); HLA-ABC FITC (W6/32; BioLegend, San Diego, CA, USA); CD19 PerCp-Cy5.5 (HIB19; BD Biosciences); anti-mouse PE-Cy5 mTer119 (TER-119; eBiosciences, San Diego, CA, USA); anti-mouse CD45.1 PE-Cy7 (A20; eBiosciences, San Diego, CA, USA); hGPA PE (HIR2; eBiosciences, San Diego, CA, USA); hCD34 APC (581; BioLegend, San Diego, CA, USA); and CD10 APC-Cy7 (HI10a; BioLegend, San Diego, CA, USA). Multi-lineage engraftment was established by the presence of myeloid cells (CD33^+^) and B-cells (CD19^+^) of engrafted human cells (CD45^+^; HLA-A/B/C^+^ cells). For GFP-expressing cells, HLA-FITC was not used in the cocktail. For secondary transplantation, only a portion of the primary mouse mononuclear population was stained, and the rest (2.5×10^5^ cells-1.3×10^6^ cells) were transplanted into six-to eight-week-old NSG mice post-irradiation conditioning. Cells were the assessed in same aforementioned manner 16wks post-transplantation into secondary mice.

### Statistical analysis

All statistical tests on experimental groups were done using Prism7 GraphPad Software. The exact statistical tests used for each comparison are noted in the figure legends.

